# cDC1 Coordinate Innate and Adaptive Responses in the Omentum required for T cell Priming and Memory

**DOI:** 10.1101/2020.07.21.214809

**Authors:** David A. Christian, Thomas A. Adams, Tony E. Smith, Lindsey A. Shallberg, Derek J. Theisen, Anthony T. Phan, Mosana Abraha, Joseph Perry, Gordon Ruthel, Joseph T. Clark, Kenneth M. Murphy, Ross M. Kedl, Christopher A. Hunter

## Abstract

The omentum in the peritoneal cavity contains fat associated lymphoid clusters (FALCs) whose role in the response to microbial challenge are poorly understood. After intraperitoneal immunization with *Toxoplasma gondii*, type I dendritic cells (cDC1) were critical to induce innate sources of IFN-γ required to recruit monocytes to the FALCs. The migration of infected peritoneal macrophages into T and B cell rich areas of the FALCs allowed the TCR-induced activation of parasite-specific T cells. Unexpectedly, cDC1 were not required for T cell priming but rather supported the expansion of parasite-specific CD8^+^ T cells. An agent-based mathematical model predicted that the lack of cDC1 would impact the early proliferative burst, and we confirmed that cDC1 were required for optimal T cell expression of nutrient uptake receptors and cell survival. These studies highlight that cDC1 in the FALCs have distinct roles in the co-ordination of the innate and adaptive responses to microbial challenge.

## INTRODUCTION

There is an increased appreciation that leukocytes present in white adipose tissue (WAT) are involved in the regulation of tissue homeostasis and the primary immune response to infection, and that this site provides a niche for immunological memory (*1*-*4*). While immune cells are present throughout the WAT, clusters of these cells form Fat Associated Lymphoid Clusters (FALCs) that densely populate the WAT tissue in the serous cavities (*5, 6*). These FALCs are positioned to provide protective immunity following a breach in porous lymphatics (*7, 8*) or the mucosal barriers in the lungs or gut (*9, 10*). In response to inflammatory or infectious challenge within serous cavities, the FALCs increase in size as leukocytes from the blood or resident cells in the cavities access these structures (*6, 10*-*12*). Compared to secondary lymphoid organs (SLOs), less is known about how FALCs are organized, how their cellular composition is altered in response to inflammatory stimuli, and whether these sites are involved in T cell priming in response to microbial challenge (*13*). The omentum is an adipose tissue present in the peritoneum that is densely populated with highly vascularized FALCs, originally termed milky spots (MS), and is a major portal for the entry of inflammatory cells into the peritoneum (*10, 11, 14, 15*). This tissue is also the site of cellular egress and antigen drainage from the peritoneal cavity (*11, 12, 14*-*19*), where the stromal cells in the MS play a role in the recruitment of monocytes and neutrophils (*15, 19*). In naïve mice, MS are dominated by a large population of B cells while also containing T cells, innate lymphoid cells (ILCs), macrophages, and conventional dendritic cells (cDCs), but these lymphoid aggregates are less organized compared to conventional SLOs (*12, 13*). Within lymph nodes (LNs) the positioning of cDC populations is critical for the efficient capture and presentation of antigen, which underlies the stimulation of protective T cell responses (*20*-*25*). At present, it is known that cDC1 reside in the WAT and play a role in whole body energy homeostasis (*26*), but little is known about the localization of cDC1 within FALCs or their role in the initiation of the innate and adaptive immune response associated with microbial challenge at serosal surfaces.

Intraperitoneal (i.p.) infection with live, invasion-competent, replication-deficient strains of *T. gondii* provide a model for vaccine-induced immune responses (*27*-*30*), and results in a parasite-specific CD8^+^ and CD4^+^ T cell response that is dependent on cDCs (*31*-*34*). While the protective T cell response to this challenge is typically assessed in the spleen, where this response is initiated is not understood. In the present study, a replication-deficient strain of *T. gondii* that expresses the model antigen ovalbumin (OVA) and Cre recombinase was combined with a Cre-sensitive reporter mouse, T cell receptor (TCR) transgenics, reporters of TCR stimulation, and a novel cDC1 reporter mouse strain to visualize the earliest events involved in T cell priming. These studies reveal that infected peritoneal macrophages migrate into distinct T and B cell regions within the MS of the omentum. *Batf3*-dependent cDC1 and IL-12 are critical to initiate the production of IFN-*γ* by ILCs that is required to recruit inflammatory monocytes and to generate the parasite-specific CD4^+^ T cell response. Canonically, the ability of cDC1 to cross present antigen is required for CD8^+^ T cell activation (*35, 36*), and we found that after CPS immunization CD8^+^ T cell priming occurs in the cDC1-enriched T cell zones of the MS. Surprisingly, in the absence of cDC1 infected macrophages still migrated to the T cell zones in the MS, and priming of CD8^+^ T cells appeared normal. However, the use of K-function statistics indicated that the lack of cDC1 resulted in a defect in the clustering of CD8^+^ T cells in the MS, and the CD8^+^ T cell response was not sustained. The use of these data sets to develop a novel agent-based model of T cell priming and clonal burst indicated that cDC1 are not required for initial priming but function to support the blastogenesis of parasite-specific CD8^+^ T cells. Consistent with this model, in *Batf3*^-/-^ mice the proliferative response of the CD8^+^ T cells was characterized by defects in nutrient receptor expression and increased cell death. Thus, cDC1 have distinct roles in co-ordination of the innate and adaptive responses in the FALCs required to generate protective CD8^+^ T cell memory.

## RESULTS

### Immunization leads to reorganization of the Omental MS

The omentum is one of the main drainage sites for interstitial fluid from the peritoneum, and after i.p. injection foreign antigens accumulate in this site (*14*). To track the distribution of the CPS strain of *T. gondii* from the peritoneum, parasites that express mCherry and inject Cre recombinase into host cells (CPS-Cre-mCherry) were used in combination with the Cre-sensitive Ai6 reporter mice to identify cells that had phagocytosed mCherry^+^ parasites (Toxo^+^ZsGreen^-^), had been infected (Toxo^+^ZsGreen1^+^), or had been injected with parasite proteins without being infected (Toxo^-^ZsGreen^+^) (*37*-*39*). Analysis of the omentum and draining parathymic and mediastinal lymph nodes (DLN) revealed that by 1 day post immunization (dpi) infected cells had moved from the peritoneum to the omentum with further accumulation by 2 dpi, while almost no infected cells were present in the DLN (Figure 1A). UMAP analysis illustrated the alteration in cellularity of the omentum from naïve and infected mice and the marked influx of large peritoneal macrophages (LPM), MHCII^+^ monocytes, and macrophages (M*ϕ*) by 2 dpi (Figure 1B, C). This analysis revealed that most interactions with the parasite occurred with CD64^+^ cells (Figure 1D). As infected cells and not cells that have phagocytosed the parasite are essential for the T cell responses to CPS (*33*), examination of which cells were productively infected or injected by the parasite indicated that LPM and M*ϕ* made up the majority of both infected cells and injected cells (Figure 1E). Because the CPS strain invades but cannot disseminate by replication and host cell lysis, these data indicate that following i.p. injection, CPS parasites primarily infect the resident macrophages that migrate into the omentum with other inflammatory cells.

**Figure 1.**
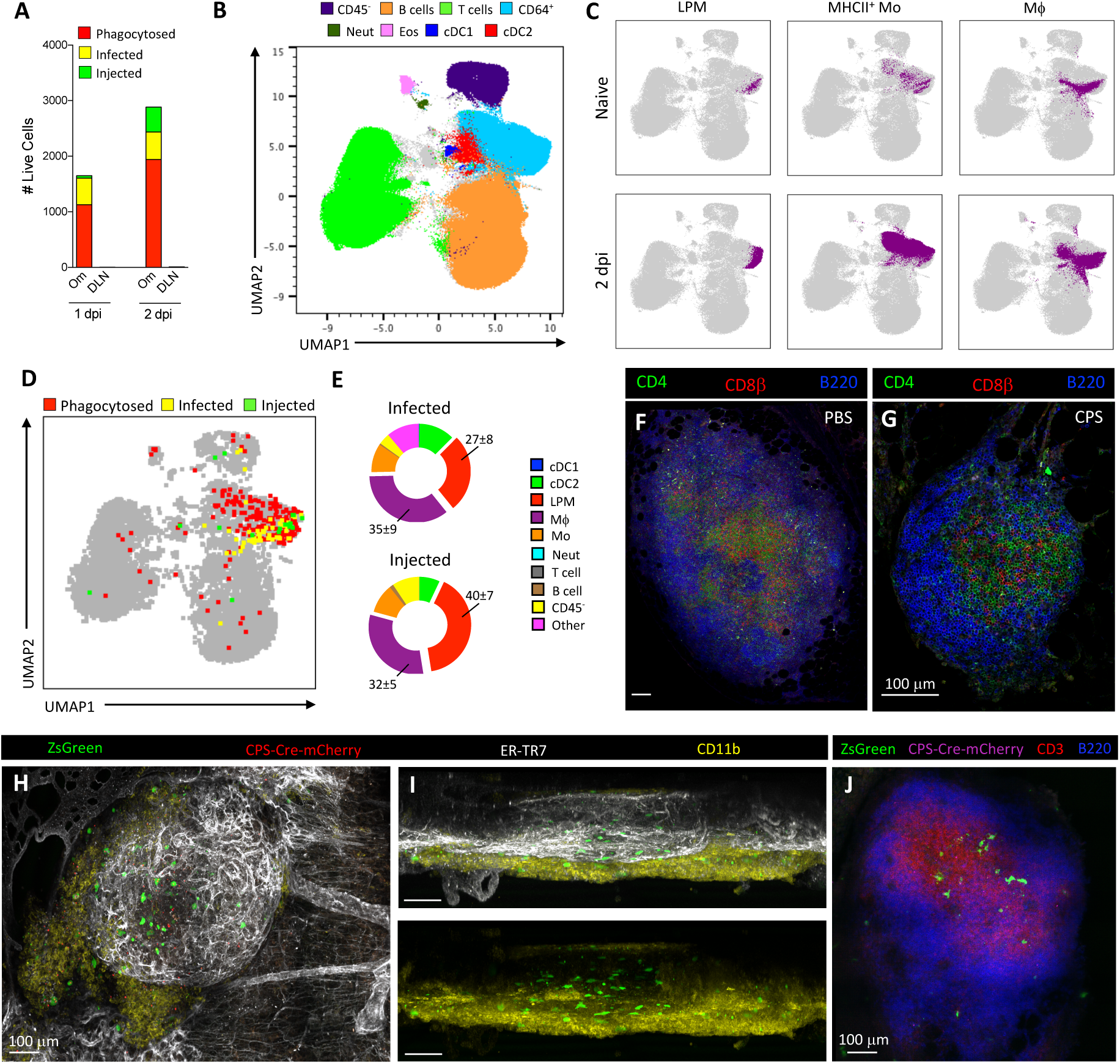
Intraperitoneal CPS immunization results in antigen transport to dynamic MS. A) Intraperitoneal immunization of Ai6 mice with 10^5^ CPS-Cre-mCherry parasites allows tracking drainage of cells phagocytosing the parasite (Toxo^+^ZsGreen^-^), infected cells (Toxo^+^ZsGreen^+^), and injected cells (Toxo^-^ZsGreen^+^) from the peritoneum to the omentum (Om) and draining mediastinal and parathymic LNs (DLN) at 1 and 2 dpi. Data (n = 4-5) represents summation of the means of each phenotype for 1 of 2 independent experiments. (B-E) UMAP analysis of cellularity of the omentum of naïve and CPS-Cre-mCherry-immunized (10^5^ parasites) Ai6 mice at 2 dpi. B) Samples show distinct clusters of non-hematopoietic cells (CD45^-^), B cells (CD45^+^CD19^+^MHCII^+^B220^+^), T cells (CD45^+^TCRβ^+^CD5^+^CD3^+^), CD64^+^ (CD45^+^CD64^+^CD11b^+^), Neut (CD45^+^CD64^-^Ly6G^+^), Eos (CD45^+^CD11b^+^SiglecF^+^), cDC1 (CD45^+^CD64^-^CD11c^+^MHCII^+^XCR1^+^CD11b^-^), cDC2 (CD45^+^CD64^-^CD11c^+^MHCII^+^XCR1^-^CD11b^+^). C)UMAP analysis of changes in the CD64^+^ population between naïve and immunized (2 dpi) Ai6 mice. LPM (CD45^+^CD64^+^CD11b^+^CD102^+^), MHCII^+^ Mo (CD45^+^CD64^+^CD11b^+^CD102^-^MHCII^+^Ly6C^+^), Mϕ (CD45^+^CD64^+^CD11b^+^CD102^+^MHCII^+^Ly6C^-^). D) UMAP analysis of cells phagocytosing parasites, infected cells, and injected cells in the omentum. E)Donut charts represent the cellular distribution of infected and injected cells shown in the UMAP analysis. Data (n = 4-6) represent 1 of 2 independent experiments. UMAP, Uniform Manifold Approximation and Projection; Neut, Neutrophil; Eos, eosinophils; cDC, conventional dendritic cell; LPM, large peritoneal macrophage; Mo, monocyte; Mϕ, macrophage. (F-G) Immunofluorescence staining of omentum MS from WT mice injected with PBS or immunized with 10^5^ CPS parasites at 2 dpi showing regions enriched in B cells and T cells. CD4 (green), CD8β (red), B220 (blue). (H-J) Immunofluorescence staining of omentum MS from Ai6 mice immunized with 10^5^ CPS-Cre-mCherry parasites at 2 dpi. (H and I) ZsGreen (green), CPS-Cre-mCherry (red), ER-TR7 (white), CD11b (yellow). (J) ZsGreen (green), CPS-Cre-mCherry (magenta), CD3 (red), B220 (blue). Scale bars in all images are 100 μm.

While the MS contains large numbers of B cells, these lymphoid aggregates have been characterized as lacking the organization typical of SLOs (*13*). Surprisingly, when the MS from PBS-injected and CPS-immunized mice were visualized as whole mount omental tissue, analysis revealed distinct regions that were enriched for B cells and a central zone that contained CD4^+^ and CD8^+^ T cells (Figure 1F-G). Next, Ai6 mice immunized with CPS-Cre-mCherry were used to visualize the localization of infected cells in the MS at 2 dpi. A stain for ER-TR7 to identify reticular cells and reticular fibers revealed an extensive glomerular vascular network, and projected images revealed that infected cells were localized in a mantle of CD11b^+^ myeloid cells (Figure 1H). Based on the use of the *lyz2*^Ai6^ reporter mouse, this mantle was largely composed of monocytes infiltrating after CPS immunization (Figure S1A-F). This projected image viewed along the XZ axis of the MS also showed that ZsGreen^+^ macrophages were dispersed throughout the vascularized MS in regions that were enriched with T and B cells (Figure 1I-J). Thus, CPS immunization induces a rapid recruitment of monocytes into the omentum and the migration of infected macrophages from the peritoneum into the distinct regions of the MS.

### cDC1 co-ordinate innate changes in the MS

cDC1 are a critical source of IL-12 that stimulates type I ILC (NK cells and ILC1s) to produce IFN-*γ* required for resistance to *T. gondii* (*40*-*46*). To determine the impact of cDC1 and IL-12 on the immune response to CPS immunization, *Batf3*^-/-^ mice that lack cDC1 in the peritoneum and omentum (Figure S2A-B) and *Il12b*^-/-^ mice were compared to WT mice. At 2 dpi whole mount imaging and flow cytometric analysis of the omentum and peritoneum revealed that immunization of WT mice resulted in an influx of monocytes, but the *Batf3*^-/-^ mice and *Il12b*^-/-^ mice had a major defect in recruitment of Ly6C^HI^ monocytes (Figure 2A-C, Figure S2C-F). Intracellular staining for IFN-*γ* at 1 dpi showed that NK cells and ILC1s produced IFN-*γ* in WT but not in *Batf3*^-/-^ or *Il12b*^-/-^ mice (Figure 2D-F, Figure S2G-I). Moreover, comparison of immunized Rag2^-/-^ and Rag2^-/-^ *γ*c^-/-^ mice (that lack these innate sources of IFN-*γ*) revealed that ILCs are required to recruit monocytes to the peritoneum (Figure S2J-K). Thus, in response to CPS, cDC1 and IL-12 are critical to stimulate ILCs that co-ordinate the recruitment of monocytes into the omentum and peritoneum. CPS immunization induces macrophage migration to the omentum and the number of LPM in the peritoneum decreases, but this was not observed in immunized *Batf3*^-/-^ and *Il12b*^-/-^ mice (Figure S2L-M). To determine if this also affects the migration of infected cells from the peritoneum to the omentum, Ai6 mice and Ai6/*Batf3*^*-/-*^ mice were immunized with CPS-Cre-mCherry. Flow cytometric analysis of infected cells in the peritoneum showed that there was a significantly higher number of total infected cells and infected LPM retained in the peritoneum of the Ai6/*Batf3*^-/-^ mice (Figure 2G). However, an equivalent number of infected and injected cells accumulated in the omentum of Ai6 and Ai6/*Batf3*^-/-^ mice at 2 dpi (Figure 2H). Analysis of the cellularity of injected or infected cells between Ai6 and Ai6/*Batf3*^-/-^ mice indicated these compositions were comparable (Figure 2I), while imaging confirmed infected macrophages were still able to migrate to the MS (Figure 2J) and into the T cell zone (Figure 2K) in Ai6/*Batf3*^-/-^ mice. Thus, while cDC1-dependent inflammation is important for the mobilization of macrophages from the peritoneum into the MS, it is not essential for migration of infected cells into the omentum.

**Figure 2.**
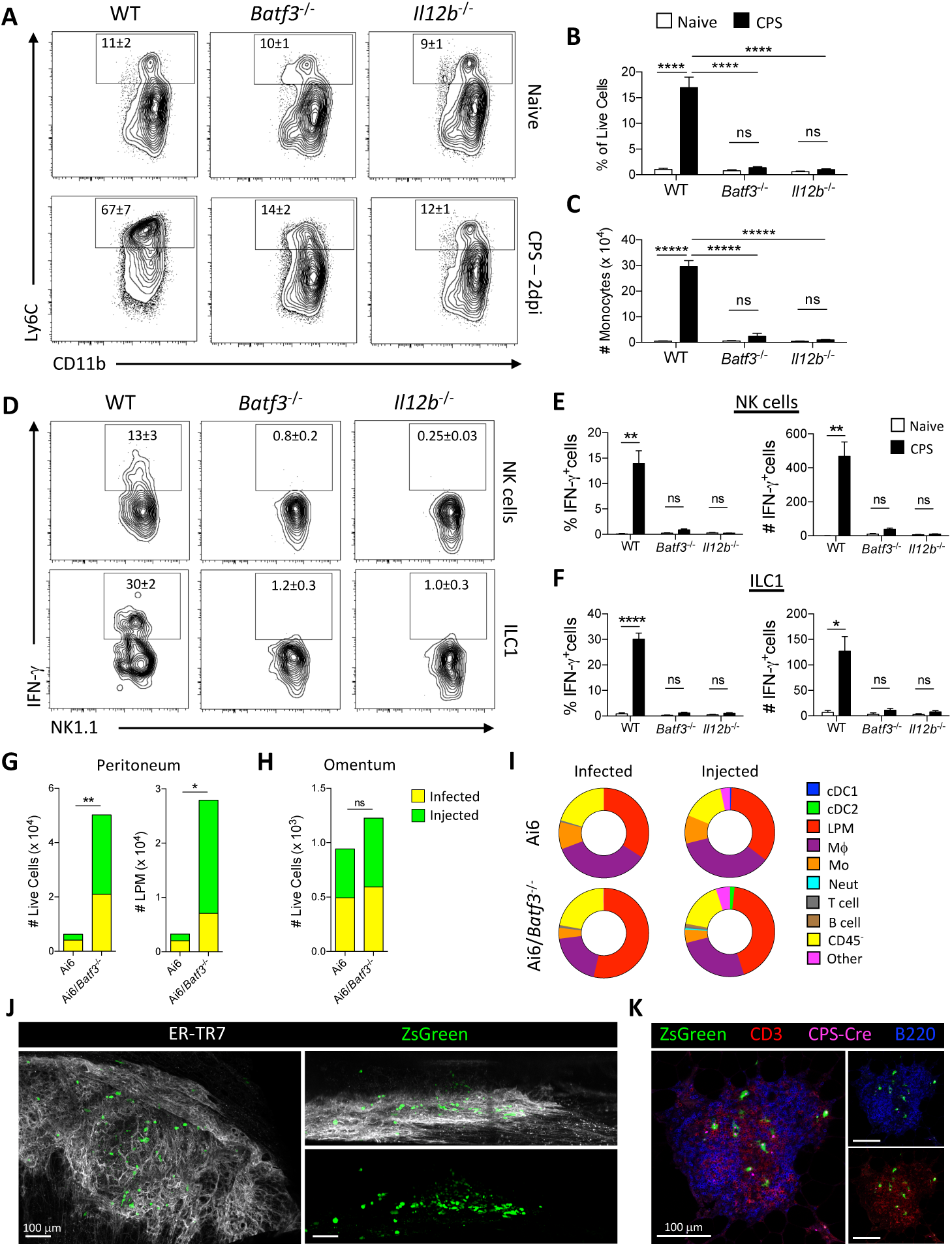
Recruitment of innate immune cells to MS requires cDC1 and IL-12. (A-C) Quantification of recruitment of monocytes to the omenta of WT, *Batf3*^-/-^, and *Il12b*^-/-^ mice after immunization with 10^5^ CPS parasites (2 dpi). (A) Flow cytometry analysis of monocytes (CD64^+^CD11b^+^CD102^-^Ly6C^HI^) in the omentum. (B and C) Quantification of monocytes (B) as a fraction of live cells in the omentum and (C) the total number of monocytes in the omentum. Data (n = 4-5) represent 1 experiment out of 2-3 independent experiments. (D-F) Quantification of IFN-γ production by Type I ILCs in the omenta of WT, *Batf3*^-/-^, and *Il12b*^-/-^ mice after immunization with 10^5^ CPS parasites (1 dpi). (D) Flow cytometry analysis of NK cells (Lin^-^NK1.1^+^NKp46^+^EOMES^+^CD200R^-^) and ILC1s (Lin^-^NK1.1^+^NKp46^+^EOMES^-^ CD200R^+^) producing IFN-γ. (E and F) Quantification comparing the fraction and total number of (E) NK cells and (F) ILC1s producing IFN-γ between naive and CPS immunized (1 dpi) mice. Lin (CD3, CD5, CD19, B220, F4/80). Data (n = 4-5) represent 1 experiment out of 2-3 independent experiments. (G-I) Quantification of infected and injected cells in peritoneum and omentum of Ai6 and Ai6/*Batf3*^-/-^ mice after immunization with 10^5^ CPS-Cre-mCherry parasites (2 dpi). (G) Number of infected and injected Live Cells and LPM in the peritoneum. (H)Number of infected and injected Live Cells in the omentum. (I)Donut charts represent the cellular distribution of infected and injected cells in the omenta of Ai6 and Ai6/*Batf3*^-/-^ mice with same gating strategy as (Fig. 1B). Data (n = 4-5) represent 1 out of 2 independent experiments. (J and K) Immunofluorescence imaging of infected cells in the MS of Ai6/*Batf3*^-/-^ mice immunized with 10^5^ CPS-Cre-mCherry parasites (2 dpi). (J) 3D rendering of infected cells entering MS. ER-TR7 (gray), ZsGreen (green). (K)Z slice of infected cells entering T cell enriched region of MS. ZsGreen (green), CD3 (red), CPS-Cre-mCherry (magenta), B220 (blue). All scale bars are 100 μm. All representative plots indicate mean ± SD. All statistical comparisons were unpaired Student’s t test. ns, not significant; *p < 0.05; **p < 0.01; ****p < 0.0001; *****p < 0.00001.

### T cell activation occurs in the omentum

To determine the impact of cDC1 on T cell responses after CPS immunization, the location of T cell priming and expansion had to be established. Therefore, CPS parasites that expressed OVA (CPS-OVA) were combined with K^b^/OVA-specific OT-I TCR transgenic mice that expressed Nur77^GFP^ (OT-I/Nur77^GFP^). These TCR transgenic T cells transiently upregulate GFP after TCR engagement by cells presenting OVA peptide via H-2K^b^ (*47*). For these experiments, OT-I/Nur77^GFP^ T cells were transferred i.p. into congenic hosts and two hours later mice were immunized with CPS or CPS-OVA parasites, and T cell responses in the peritoneum and omentum were examined at 18 hours post-immunization (hpi). The combination of upregulation of CD69 combined with expression of Nur77^GFP^ was used as a measure of recent TCR engagement. In the omentum of mice that received CPS, 3±3% of OT-I/Nur77^GFP^ T cells were CD69^high^ and Nur77^GFP+^, whereas in mice immunized with CPS-OVA 43±3% of these T cells were CD69^high^Nur77^GFP+^ (Figure 3A-B). The number of activated CD69^high^Nur77^GFP+^ OT-I T cells was significantly increased in the omentum by 2 dpi compared to the DLN (Figure 3C), implicating the omentum as the site of CD8^+^ T cell priming. Next, OT-I/Nur77^GFP^ T cells labeled with CellTrace Violet (CTV) were transferred into WT mice that were then immunized with CPS-OVA and the T cell response at 2, 3, 4, 6, and 8 dpi in the omentum, DLN, and spleen was assessed (Figure 3D). At 2 dpi in the omentum 31±9% of OT-I T cells were Nur77^GFP+^ and started to divide as measured by CTV dilution (CTV^diluted^), and few OT-I T cells were detected in the DLN and spleen (Figure 3D). By 3 and 4 dpi the OT-I T cells in the omentum had undergone robust proliferation and significant expansion (89±16% and 96±3% Nur77^GFP+^/CTV^diluted^, respectively), while >90% of OT-I T cells in the DLN and spleen at 4 dpi had divided 5 or more times. At later time points (6 and 8 dpi) OT-I T cells that had completely diluted the CTV and were readily observed in the omentum, DLN, and spleen.

**Figure 3.**
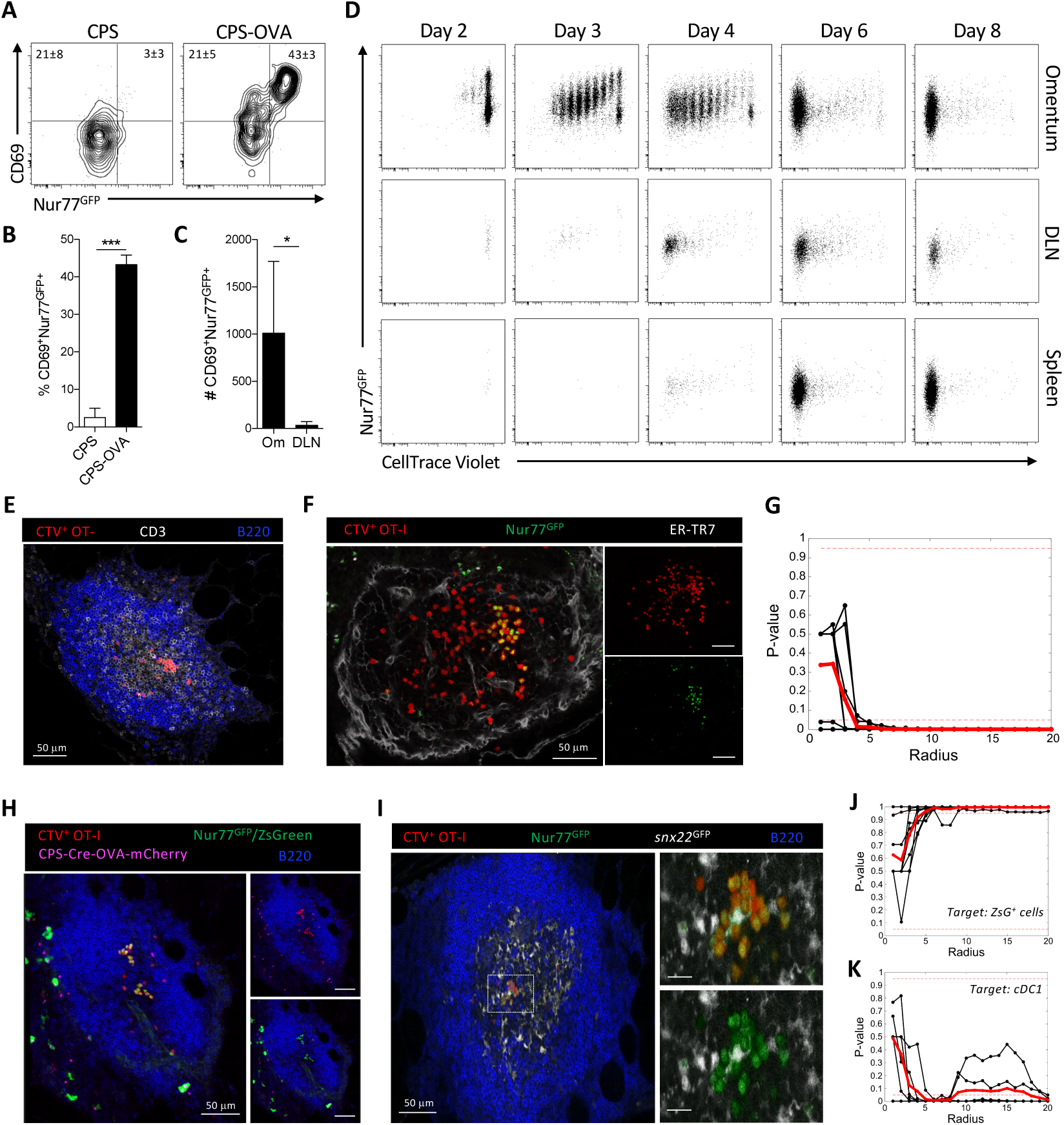
CD8^+^ T cell activation in the omentum. (A and B) Intraperitoneal transfer of 10^6^ CTV-labeled OT-I/Nur77^GFP^ T cells into WT mice that were then immunized with 2 × 10^5^ CPS or CPS-OVA parasites 2 hr later. (A) Flow cytometric analysis of activation of transferred CTV^+^ OT-I/Nur77^GFP^ T cells in the omentum of recipient WT mice at 18 hpi measured by expression of CD69 and Nur77^GFP^. (B)Quantification of the fraction of activated (CD69^+^Nur77^GFP+^) OT-I/Nur77^GFP^ T cells in the omentum. Data (n = 4-6) represent 1 experiment out of 3 independent experiments. (C) Intraperitoneal transfer of 10^6^ congenic OT-I/Nur77^GFP^ T cells into WT mice immunized with 2 × 10^5^ CPS-OVA parasites 2 hr later. Quantification of the number of activated (CD69^+^Nur77^GFP+^) OT-I/Nur77GFP T cells in the Om and DLN at 18 hpi. Data (n = 4-6) represent 1 experiment out of 2 independent experiments. (D-G) Intraperitoneal transfer of 5 × 10^5^ CTV-labeled OT-I/Nur77^GFP^ T cells into WT mice immunized with 2 × 10^5^ CPS-OVA parasites 2 hr later. (D) Representative flow cytometric analysis of the activation (Nur77^GFP+^) and division (CTV dilution) of OT-I T cells in the omentum, DLN, and spleen at 2, 3, 4, 6, and 8 dpi. Data (n = 3-5) represent 1 experiment out of 2 independent experiments. (E-G) Imaging and analysis of OT-I/Nur77^GFP^ T cell clustering in the MS of WT mice immunized with CPS-OVA at 18 hpi. (E) Immunofluorescence imaging of OT-I T cell clustering in the T cell enriched area of the MS. CTV (red), CD3 (gray), B220 (blue). Scale bar is 50 μm. (F) Immunofluorescence imaging of CTV^+^ OT-I/Nur77^GFP+^ T cell clustering in MS. CTV (red), Nur77^GFP^ (green), ER-TR7 (gray). Scale bar is 50 μm. (G) K-function statistical analysis of Nur77^GFP+^ OT-I T cell clustering in the MS of WT mice immunized with CPS-OVA at 18 hpi. Analysis of clustering in 9 individual MS (black lines) from 2 individual mice was averaged (red line). Dashed red lines indicate significant P values of 0.05 and 0.95. Data represent 1 experiment out of 7 independent experiments. (H-K) Intraperitoneal transfer of 5 × 10^5^ CTV-labeled OT-I/Nur77^GFP^ T cells into Ai6 or *snx22*^GFP/+^ mice that were immunized with 2 × 10^5^ CPS-Cre-OVA-mCherry parasites 2 hr later. Imaging and analysis of OT-I/Nur77^GFP^ T cell clusters in the MS at 18 hpi. (H) Immunofluorescence imaging of infected cells and CTV^+^ OT-I/Nur77^GFP+^ T cell clustering in MS of Ai6 mice. CTV (red), Nur77^GFP^ and ZsGreen (green), CPS-Cre-OVA-mCherry (magenta), B220 (blue). Scale bars are 50 μm. (I) Immunofluorescence imaging of cDC1 and CTV^+^ OT-I/Nur77^GFP+^ T cell clustering in the MS of *snx22*^GFP/+^ mice. CTV (red), Nur77^GFP^ (green), *snx22*^GFP^ (gray), B220 (blue). Nur77^GFP^ and *snx22*^GFP^ were distinguished and pseudocolored differently using the channel subtraction application in Imaris imaging software: *snx22*^GFP^ = (total GFP) – CTV. All scale bars are 50 μm. (J-K) Cross K-function statistical analysis of Nur77^GFP+^ OT-I T cell clusters compared to the localization of target cell population. (J) Cluster localization compared to ZsGreen^+^ cells in the MS of Ai6 mice. Analysis of clustering in 9 individual MS (black lines) from 2 individual mice was averaged (red line). Dashed red lines indicate significant P values of 0.05 and 0.95. Data represent 1 experiment out of 3 independent experiments. (K) Clustering localization compared to cDC1 in the MS of *snx22*^GFP/+^ mice. Analysis of clustering in 6 individual MS (black lines) from 2 individual mice was averaged (red line). Dashed red lines indicate significant P values of 0.05 and 0.95. Data represent 1 experiment out of 3 independent experiments. CTV, CellTrace Violet; Om, Omentum; DLN, draining LN. Representative plots in (B) and (C) indicate mean ± SD. Statistical comparisons in (B) and (C) were unpaired Student’s t test. *p < 0.05; ***p < 0.001.

Whole mount imaging of the omentum at 18 hpi with CPS-OVA revealed that CTV^+^ OT-I T cells were present in the T cell zone of the MS and a portion of these were in aggregates (Fig 3E). Additional analysis revealed that these clusters contained CTV^+^Nur77^GFP+^ OT-I T cells (a hallmark of T cell priming (*48*)) while CTV^+^Nur77^GFP-^ OT-I T cells appeared to be more dispersed (Figure 3F). To determine if there were significant differences in cluster formation between these Nur77^GFP+^ and Nur77^GFP-^ OT-I T cells, random permutation tests of clustering were employed using K-function statistics (see Methods and Supplemental Information) to determine if the Nur77^GFP+^ OT-I T cells were more likely to exist in clusters compared to the total OT-I population within the same MS. In this analysis, the Nur77^GFP+^ OT-I T cells exhibited significant clustering at a radius of *k* = 4 μm (Figure 3G). Next, cross K-function statistics were used (see Methods and Supplemental Information) to determine whether infected or injected myeloid cells (identified as ZsGreen^+^ in Ai6 mice, Figure 3H) or cDC1 (*snx22*^GFP+^ (*49*), Figure 3I) were associated with activated T cells. For this evaluation, the average number of cells of either APC population within distance *k* of each Nur77^GFP+^ OT-I T cell was compared with those of randomly permuted populations. The results showed that Nur77^GFP+^ OT-I T cells exhibited no significant clustering around ZsGreen^+^ infected cells (Figure 3J) at any scale, and were significantly dispersed away from these cells at all radii larger than 5 μm. In contrast, the Nur77^GFP+^ OT-I T cells were significantly clustered around cDC1 present in the T cell zone of the MS (Supplemental Movie 1) at radii starting around 5 μm (Figure 3K). These data establish not only that uninfected cDC1 associate with CD8^+^ T cells during early T cell priming events in the MS of the omentum, but also that the early proliferation following antigen encounter by T cells occurs entirely in the omentum.

### cDC1 are required for CPS-induced T cell responses

To assess the role of cDC1 in the generation of CPS induced CD4^+^ and CD8^+^ T cell responses, WT and *Batf3*^-/-^ mice received a single i.p. vaccination with CPS parasites. Upon challenge with the lethal RH strain 30 days later, immunized WT mice were completely protected while - similar to naïve WT mice - immunized *Batf3*^-/-^ mice had increased parasite numbers (data not shown) and succumbed to the secondary challenge (Figure 4A). The susceptibility of *Batf3*^-/-^ mice after CPS immunization was characterized by a severely reduced endogenous parasite-specific (tgd057:K^b+^) CD8^+^ T cell population at 30 dpi (Figure 4B). Analysis of these tgd057:K^b+^ CD8^+^ T cells revealed a significant difference in the memory (Tmem, CXCR3^+^KLRG1^-^) and effector (T_eff_, CXCR3^-^KLRG1^+^) phenotypes (Figure 4C-D). This defect in tgd057:K^b+^ CD8^+^ T cells in the spleen was observed as early as 9 dpi (Figure 4E), and was accompanied by an impaired parasite-specific (AS15:I-A^b+^) CD4^+^ T cell response (Figure 4F). The tgd057:K^b+^ CD8^+^ T cell response was also significantly altered at the site of immunization in the peritoneum (Figure 4G), where 30±11% of these parasite-specific T cells expressed a T_eff_ phenotype in WT mice that was significantly diminished (5±1%) in the *Batf3*^-^/- mice (Figure 4H). Instead, tgd057:Kb+ CD8+ T cells had an increased proportion of Tmem cells in the *Batf3*^-/-^ mice (65±6%) compared to WT mice (28±8%). The defect in the CD8^+^ T cell response was cell-extrinsic as the i.p. transfer of *Batf3*-sufficient OT-I T cells into WT or *Batf3*^-/-^ mice immunized with CPS-OVA resulted in fewer OT-I T cells in *Batf3*^-/-^ mice at 8 dpi and a loss of the T_eff_ CD8^+^ T cell phenotype (Figure 4I-J). These data sets are consistent with a role for cDC1 in priming CD8^+^ T cells (*50, 51*), but also suggested that cDC1 are important for the activation of CD4^+^ T cells.

**Figure 4.**
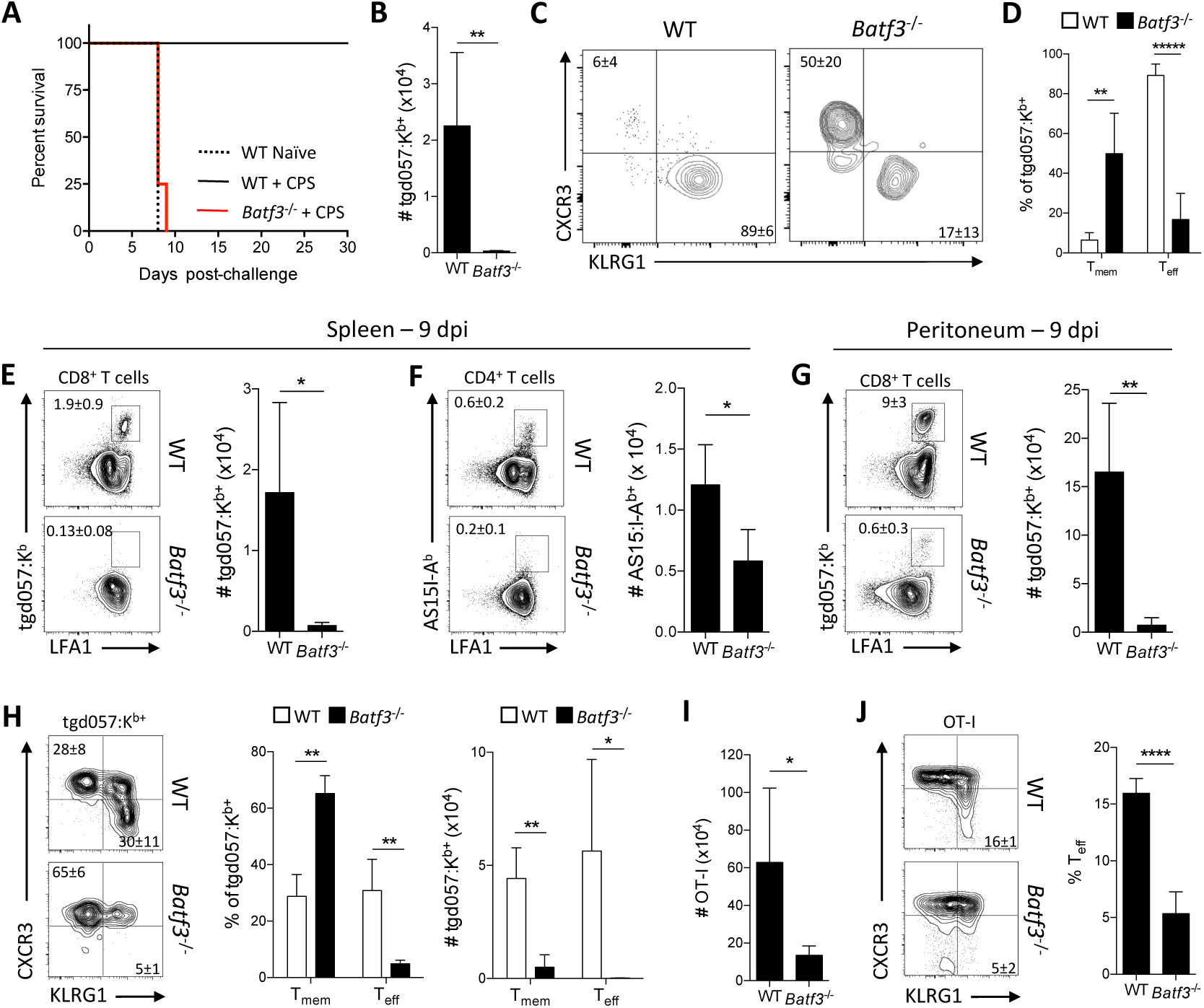
T cell responses to CPS require cDC1. (A) WT and *Batf3*^-/-^ mice were immunized with 10^5^ CPS parasites. At 30 dpi, immunized WT and *Batf3*^-/-^ mice and naïve WT mice were challenged with 10^4^ tachyzoites of the lethal RH strain of *T. gondii* and monitored for survival. A survival curve is shown. Data (n = 4-5) represent 1 experiment out of 2 independent experiments. (B-D) WT and *Batf3*^-/-^ mice were immunized with 10^5^ CPS parasites. At 30 dpi T cell populations in the spleen were evaluated by flow cytometry. (B) Number of *T. gondii*-specific (tgd057:K^b+^) CD8^+^ T cells. (C) Representative plots of the Tmem (CXCR3+KLRG1-) or T_eff_ (CXCR3-KLRG1+) phenotypes of tgd057:Kb+ CD8+ T cells. (D) The fraction of Tmem and T_eff_ phenotypes of tgd057:Kb+ CD8+ T cells. Data (n = 5) represent 1 experiment out of 2 independent experiments. (E-H) WT and *Batf3*^-/-^ mice were immunized with 10^5^ CPS parasites. At 9 dpi T cell populations were evaluated by flow cytometry. (E) Representative plots and the number of tgd057:K^b+^ CD8^+^ T cells in the spleen. (F) Representative plots and the number of *T. gondii*-specific (AS15:I-A^b+^) CD4^+^ T cells in the spleen. (G) Representative plots and the number of tgd057:K^b+^ CD8^+^ T cells in the peritoneum. (H) Representative plots and quantification of the fraction and number of Tmem or T_eff_ phenotypes of tgd057:Kb+ CD8+ T cells in the peritoneum. Data (n = 4-5) represent 1 experiment out of 3 independent experiments. (I-J) 5 × 10^5^ congenic OT-I T cells were transferred i.p. into WT and *Batf3*^-/-^ recipient mice that were then immunized with 2 × 10^5^ CPS-OVA parasites 2 hr later. OT-I T cell populations were evaluated by flow cytometry at 9 dpi in the peritoneum. (I) Number of OT-I T cells. (J) Representative plots of OT-I T cell phenotypes and fraction of T_eff_ OT-I T cells. Data (n = 4-5) represent 1 experiment out of 2 independent experiments. All representative plots indicate mean ± SD. All statistical comparisons were unpaired Student’s t test. *p < 0.05; **p < 0.01; ****p < 0.0001; *****p < 0.00001.

The ability of cDC1 to produce IL-12 is required to generate a protective immune response during toxoplasmosis (*50*). To test whether treatment of *Batf3*^-/-^ mice with exogenous IL-12 would restore the CD4^+^ and CD8^+^ T cell response, WT and *Batf3*^-/-^ mice were treated with recombinant IL-12p70 (rIL-12) during the first 3 days of CPS immunization and the T cell response in the peritoneum was assessed at 11 dpi. In WT mice, treatment with rIL-12 did not affect the AS15:I-A^b+^ CD4^+^ T cell response, but in the *Batf3*^-/-^ mice it restored the frequency of these CD4^+^ T cells to WT levels (Figure 5A-B) and increased the number compared to PBS treatment (Figure 5C). In contrast, addition of rIL-12 did not alter the CD8^+^ T cell response in WT mice, and in the *Batf3*^-/-^ mice it did not increase the frequency or number of tgd057:K^b+^ CD8^+^ T cells (Figure 5D-F) or restore the differentiation of T_eff_ CD8^+^ T cells (Figure 5G-I). These data indicate that cDC1 are an early source of IL-12 that is important for CD4^+^ T cell responses, but cDC1 have a distinct role in the induction of a protective, parasite-specific CD8^+^ T cell response.

**Figure 5.**
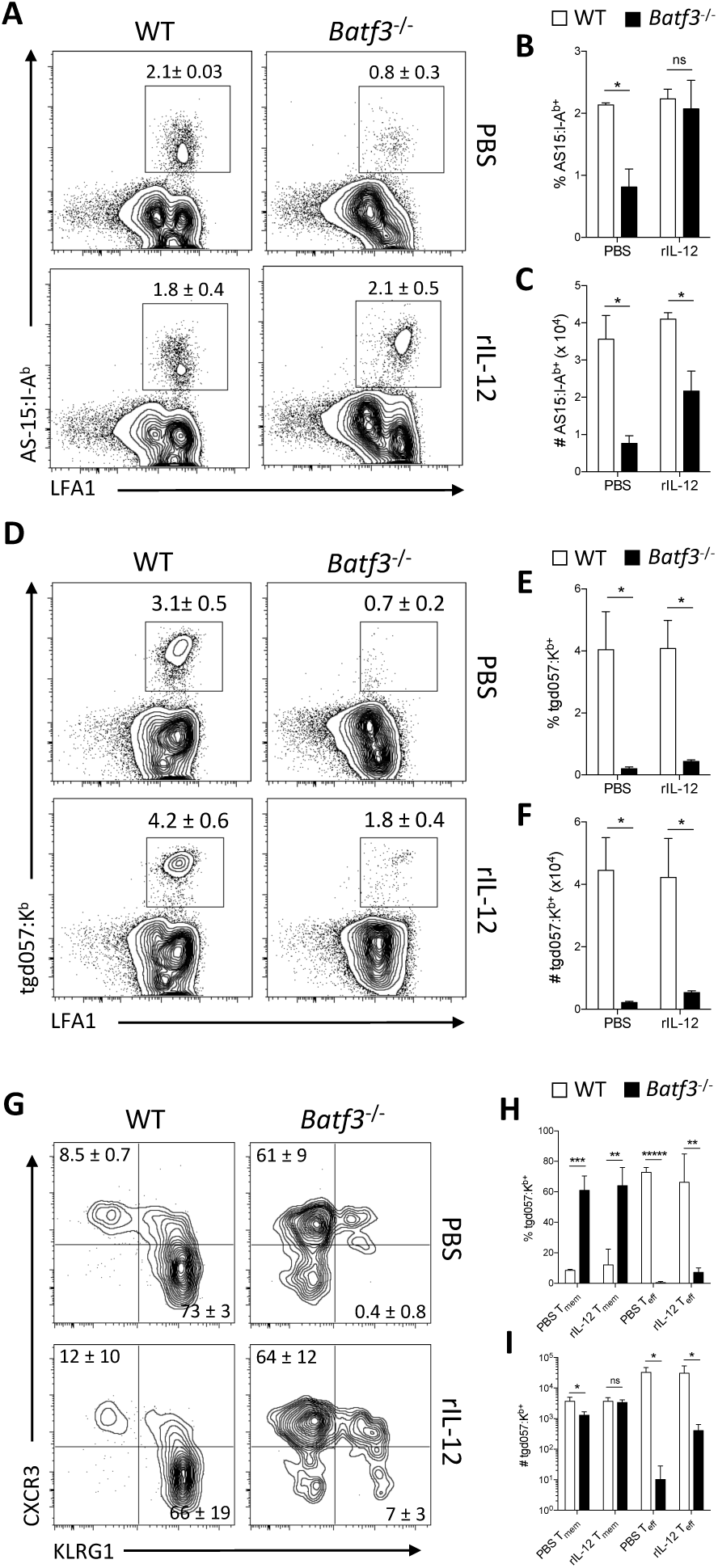
An IL-12 independent role for cDC1s in the CD8^+^ T cell response to CPS. (A-I) WT and *Batf3*^-/-^ mice were immunized with 10^5^ CPS parasites followed by i.p. administration of either PBS or 200 ng rIL-12 at 0, 24, and 48 hpi. T cell responses in the peritoneum were assessed via flow cytometry at 11 dpi. (A) Representative plots of AS15:I-A^b+^ CD4^+^ T cells. (B-C) The fraction and number of AS15:I-A^b+^ CD4^+^ T cells. (D) Representative plots of tgd057:K^b+^ CD8^+^ T cells. (E-F) The fraction and number of tgd057:K^b+^ CD8^+^ T cells. (G)Representative plots of the T_eff_ and Tmem phenotypes of tgd057:K^b+^ CD8^+^ T cells. (H-I) The fraction and number of Tmem or T_eff_ phenotypes of tgd057:K^b+^ CD8^+^ T cells. Data (n = 3-5) represent 1 experiment out of 2 independent experiments. All representative plots indicate mean ± SD. All statistical comparisons were unpaired Student’s t test. ns, not significant; *p < 0.05; **p < 0.01; ***p < 0.001; *****p < 0.00001.

### cDC1 are not required for CD8^+^ T cell priming but support T cell expansion

The pr0iming and expansion of T cells in the omentum of *Batf3*^-^/- mice was examined to define when cDC1 were required for the generation of a protective CD8^+^ T cell response. Congenic OT-I/Nur77^GFP^ T cells were transferred into WT and *Batf3*^-/-^ mice that were then immunized with CPS-OVA. Surprisingly, at 18 hpi OT-I/Nur77^GFP^ T cells in WT and *Batf3*^-/-^ mice showed a similar increase in the fraction of newly activated CD69^+^Nur77^GFP+^ OT-I T cells (Figure 6A-B). However, imaging of OT-I/Nur77^GFP^ T cell priming in the omentum of *Batf3*^-/-^ mice showed that although these T cells receive TCR stimulation by 18 hours, Nur77^GFP+^ OT-I T cells did not show significant clustering compared to Nur77^GFP-^ OT-I T cells (Figure 6C-D).

**Figure 6.**
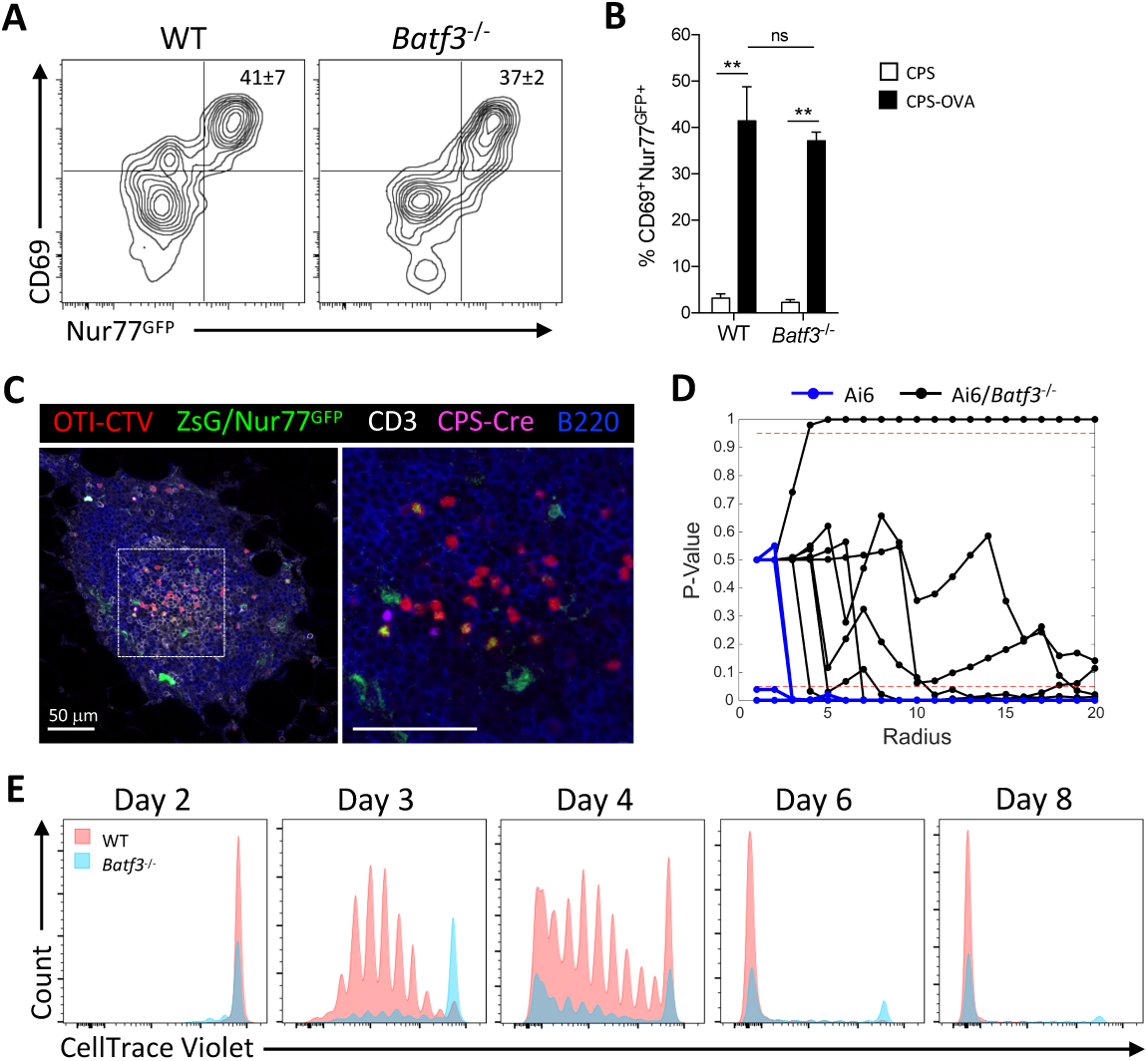
cDC1 required for CD8^+^ T cell expansion but not activation. (A and B) Intraperitoneal transfer of 10^6^ congenic OT-I/Nur77^GFP^ T cells into WT and *Batf3*^-/-^ mice that were then immunized with 2 × 10^5^ CPS or CPS-OVA parasites 2 hr later. Activation of OT-I/Nur77GFP T cells in the omentum at 18 hpi was analyzed by flow cytometry. (A) Representative plots of the activation of transferred OT-I/Nur77^GFP^ T cells measured by expression of CD69 and Nur77^GFP^. (B) Quantification of the fraction of activated (CD69^+^Nur77^GFP+^) OT-I/Nur77^GFP^ T cells in the omentum. Data (n = 3-4) represent 1 experiment out of 2 independent experiments. Unpaired Student’s t test was performed (mean ± SD). (C) Intraperitoneal transfer of 5 × 10^5^ CTV-labeled OT-I/Nur77^GFP^ T cells into Ai6/*Batf3*^-/-^ mice that were immunized with 2 × 10^5^ CPS-Cre-OVA-mCherry parasites 2 hr later. Immunofluorescence imaging of CTV^+^ OT-I/Nur77^GFP^ T cell clustering in the MS of Ai6/*Batf3*^-/-^ mice at 18 hpi. CTV (red), ZsGreen, Nur77^GFP^ (green), CD3 (gray), CPS-Cre-OVA-mCherry (magenta), B220 (blue). Scale bar is 50 mm. (D) Intraperitoneal transfer of 5 × 10^5^ CTV-labeled OT-I/Nur77^GFP^ T cells into Ai6 and Ai6/*Batf3*^-/-^ mice that were immunized with 2 × 10^5^ CPS-Cre-OVA-mCherry parasites 2 hr later. K-function statistical analysis of Nur77^GFP+^ OT-I T cell clustering in individual MS from Ai6 mice (blue lines) and Ai6/*Batf3*^-/-^ mice (black lines). Analysis of clustering in Ai6 mice was done for 9 individual MS from 2 individual mice, and analysis in Ai6/*Batf3*^-/-^ mice was done for 7 individual MS from 2 individual mice. Data represent 1 experiment out of 4 independent experiments. (E) Intraperitoneal transfer of 5 × 10^5^ CTV-labeled OT-I/Nur77^GFP^ T cells into WT and *Batf3*^-/-^ mice immunized with 2 × 10^5^ CPS-OVA parasites 2 hr later. Representative flow cytometric analysis of the division (CTV dilution) of OT-I T cells in the omentum at 2, 3, 4, 6, and 8 dpi. Data (n = 3-5) represent 1 experiment out of 2 independent experiments. WT recipient mice (red histograms), *Batf3*^-/-^ recipient mice (blue histograms). ns, not significant; **p < 0.01.

Intact OT-I T cell priming in *Batf3*^-/-^ mice at 18 hpi indicated that the defect in the OT-I T cell response in these mice occurred during the T cell expansion phase. Therefore, the kinetics of the OT-I T cell response to CPS-OVA was measured at 2, 3, 4, 6, and 8 dpi in WT and *Batf3*^-/-^ mice (Figure 6E). At 2 dpi in the omentum there continued to be no defect in the activation and early proliferative response of OT-I T cells in the *Batf3*^-/-^ mice compared to in the WT mice. However, by 3 dpi the OT-I T cells in the WT mice had undergone robust proliferation and significant expansion such that 82±21% had diluted CTV, whereas in *Batf3*^-/-^ mice only 43 ±16% OT-I T cells had diluted CTV (Figure 6E). This defect in expansion persisted through 8 dpi, which resulted in a significantly decreased number of OT-I T cells in the *Batf3*^-/-^ mice (see Figure 4I). These data indicate that in vaccinated *Batf3*^-/-^ mice there is no defect in the early activation of OT-I T cells, but the absence of cDC1 results in impaired clustering and a defect in the expansion phase of the response.

In order to understand the differences in the CD8^+^ T cell response between WT and *Batf3*^-/-^ mice, a reductionist approach was used to generate an agent-based model (STochastic Omentum Response, STORE), that performs stochastic simulations of the generation of a protective T cell response. First, a combination of set parameters that were based on literature values (Figure 7A) and six variable parameters derived from our experiments were utilized: 1) the number of transferred naïve OT-I, 2) the ratio of transferred OT-I T cells to the number of APCs generated by CPS-OVA administration (*r*_*AP*_), the probabilities of 3) an OT-I T cell and APC binding (*p*_*AP,bind*_), 4) an APC leaving the system (*p*_*BP,leave*_), 5) cells in divisions 3-6 leaving (*p*_*EH,leave*_), and 6) cells in divisions 7-8+ leaving (*p*_*IJ,leave*_). After an iterative process this stochastic model was able to track the behavior and fate of individual OT-I T cells in the omentum of each mouse for 8 days after CPS-OVA immunization. A visual demonstration of the stochasticity of the model and its fidelity is provided in Figure 7B-E in which the data points are the experimental average cell counts of OT-I T cells at 2, 3, 4, 6, and 8 dpi and the lines are an example simulation run using the STORE model. This example has a large OT-I/APC injection ratio (*rAP* = 5.1), which makes the APC population the rate limiting factor and allows naïve OT-I T cells (A) to accumulate rapidly over the first 30 hours and then decay exponentially for the remainder of the 8 day period (Figure 7B). In this simulation the number of free APCs (P) remains low throughout the simulation as any available APC almost immediately pairs with a naïve OT-I T cell (AP) (Figure 7C) and the time for GFP expression and conversion to a Nur77^GFP+^ OT-I:APC pair (BP) is roughly 30 minutes. The duration of the BP population is roughly 24 hours to approximate the second phase of DC:T cell interactions during T cell priming (*48*). The BP trajectory features a linear reduction toward zero because the model approximates a linear decrease in the APC population over time (*p*_*BP,leave*_). Simulation of divisions 1-6 (Figure 7D-E) reflect the 24 hour T cell priming cycle of BP and the 5.3 hour division cycle (*52*) which is apparent in the approximately 5.3 hour phase difference in cycles between divisions. The effect of the linear reduction of BP over time (and thus the number of primed T cells that begin to divide) is evidenced by the decreasing cell counts in divisions 1-6 over time (Figure 7D-E). Division 8+ (J) has a different profile (Figure 7E), because these cells are all CTV^-^ and thus continued division of these cells (*53*) results in a growth trajectory that is exponential at first and then flattens as the number of precursor cells begins to disappear by division or by exiting the omentum. Simulations of the kinetics of the OT-I T cell response of WT and *Batf3*^-/-^ mice using the optimized STORE model were then performed and compared to data from two experiments which examined the kinetics of the OT-I T cell response, measured by Nur77^GFP^ or CD69 expression and division rate by dilution of CTV, in WT and *Batf3*^-/-^ recipient mice (Figure 7B-E). Simulation results for the number of each OT-I T cell type matched the experimental data at each time point for both WT and *Batf3*^-/-^ mice reasonably well and further validated the model (Figure 7F). This comparison highlighted that the most significant difference between WT and *Batf3*^-/-^ mice is the probability of OT-I T cells in divisions 3-6 leaving the system by 68 hpi. This difference could be due to migration out of the omentum or a defect in cell survival or blastogenesis. As there was not an increased number of OT-I T cells in the DLN or spleen of *Batf3*^-/-^ mice compared to WT mice (data not shown), OT-I T cells in divisions 3-6 were examined for differing rates of cell death in WT and *Batf3*^-/-^ mice at 3 dpi. OT-I T cells isolated from the omentum were stained for terminal cell death via Annexin V and propidium iodide (PI), which showed increased cell death in divisions 3 and 4 in *Batf3*^-/-^ mice compared to WT mice (Figure 7G-H).

**Figure 7.**
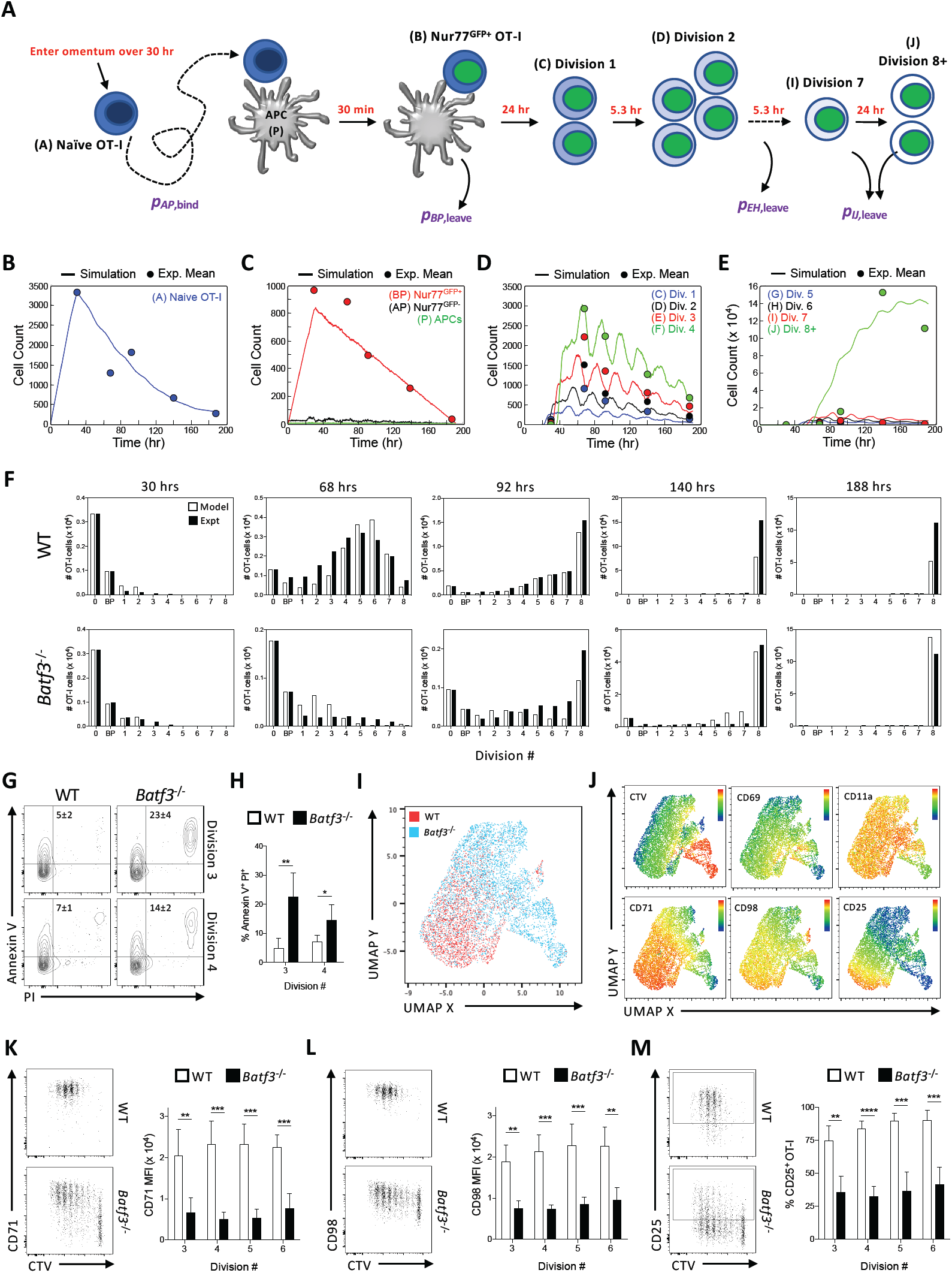
Stochastic modeling implicates defect in OT-I expansion in *Batf3*^-/-^ mice. (A) Schematic of STORE stochastic model of OT-I T cell activation and expansion. Set parameters (red text) in the model included 1) entry of naïve OT-I T cells and APCs at a linear rate over 30 hr (Δ*t*_*arrival*_), 2) maturation time of Nur77GFP at 30 min, 3) activated OT-I T cells clustering on an APC for 24 hr, 4) 5.3 hr division rate of activated OT-I T cells in divisions 1-6, 5) 24 hr division rate for OT-I T cells beyond division 6. The six variable parameters (purple text) included 1) the number of transferred naïve OT-I, 2) the ratio of transferred OT-I T cells to the number of APCs generated by CPS-OVA administration (*r*_*AP*_), the probabilities of 3) an OT-I T cell and APC binding (*p*_*AP,bind*_), 4) an APC leaving the system (*p*_*BP,leave*_), 5) cells in divisions 3-6 leaving (*p*_*EH,leave*_), and 6) cells in divisions 7-8 leaving (*p*_*IJ,leave*_. (B-F) Intraperitoneal transfer of 5 × 10^5^ OT-I/Nur77^GFP^ T cells into WT and *Batf3*^-/-^ mice immunized with 2 × 10^5^ CPS-OVA parasites 2 hr later. OT-I T cells in each division in the omentum at 2, 3, 4, 6, and 8 dpi were quantified by flow cytometry. (B-E) Representative plots from a simulation of the STORE model where solid lines indicate simulation data and filled circles indicate experimental data. (B) The number of naïve OT-I T cells over time. (C) The numbers of unbound APCs and APC binding pairs with Nur77^GFP-^ (AP) and Nur77^GFP+^ (BP) OT-I T cells over time. (D) The number OT-I T cells in divisions 1-4 versus time. (E)The number of OT-I T cells in divisions 5-8+ versus time. (F) The number of OT-I T cells at different time points from model simulations and experimental data in both WT and *Batf3*^-/-^recipient mice. Data (n = 4-5) represent 1 experiment out of 2 independent experiments. (G-H) Intraperitoneal transfer of 5 × 10^5^ CTV-labeled OT-I/Nur77^GFP^ T cells into WT and *Batf3*^-/-^ mice immunized with 2 × 10^5^ CPS-OVA parasites 2 hr later. Cell death of OT-I T cells was measured in the omentum at 3 dpi. (G) Representative plots of measured cell death (Annexin V^+^ PI^+^) in OT-I T cells in divisions 3 and 4 in WT and *Batf3*^-/-^ mice. (H) The fraction of dead OT-I T cells in divisions 3 and 4 in WT and *Batf3*^-/-^ mice. Data (n = 5) represent 1 experiment out of 2 independent experiments. (I-M) Intraperitoneal transfer of 5 × 10^5^ CTV-labeled OT-I/Nur77^GFP^ T cells into WT and *Batf3*^-/-^ mice immunized with 2 × 10^5^ CPS-OVA parasites 2 hr later. OT-I T cells in the omentum at 3 dpi were evaluated by flow cytometry. (I) UMAP analysis showing OT-I T cells from WT and *Batf3*^-/-^ mice. (J) UMAP analysis of the intensity of CTV and expression levels of CD69, CD11a, CD71, CD98, and CD25 by OT-I T cells from WT and *Batf3*^-/-^ mice. (K) Representative plot of the expression of CD71, and quantification of the MFI of CD71 expression by divisions 3-6 of OT-I T cells from WT and *Batf3*^-/-^ mice. (L) Representative plot of the expression of CD98, and quantification of the MFI of CD98 expression by divisions 3-6 of OT-I T cells from WT and *Batf3*^-/-^ mice. (M) Representative plot of the expression of CD25, and quantification of the fraction of CD25^+^ OT-I T cells in divisions 3-6 of OT-I T cells from WT and *Batf3*^-/-^ mice. Data (n = 4-5) represent 1 experiment out of 6 independent experiments. STORE, STochastic Omentum REsponse; PI, propidium iodide. All representative plots indicate mean ± SD. All statistical comparisons were unpaired Student’s t test. *p < 0.05; **p < 0.01; ***p < 0.001; ****p < 0.0001.

The blastogenic stage of the T cell response requires increased metabolic activity driven by aerobic glycolysis and mitochondrial activity, an enhanced responsiveness to IL-2, and nutrient uptake required for rapid division. OT-I T cells transferred into WT and *Batf3*^-/-^ mice were first examined at 3 dpi for changes in mitochondria and glucose uptake. In OT-I T cells that had become activated (CD69^+^) and begun blastogenesis in either WT or *Batf3*^-/-^ mice, there was no difference in the mitochondrial mass, surface expression of the glucose uptake receptor Glut1, or glucose uptake (Figure S3A-C). UMAP analysis of expression of a panel of T cell activation-dependent receptors and CTV dilution at 3 dpi differentiated between OT-I T cells from WT versus *Batf3*^-/-^ recipients (Figure 7I). Heatmap overlays of the UMAP plot for CTV dilution and expression levels of markers of T cell activation (CD69 and CD11a) showed little difference between OT-I T cells from WT versus *Batf3*^-/-^ recipients (Figure 7J). However, heatmap overlays for the expression of the nutrient uptake receptors, CD71 and CD98, as well as the cytokine receptor IL-2R*α* (CD25) showed increased expression of all three receptors in OT-I T cells from WT recipients (Figure 7J). The expression level of CD71, CD98, and CD25 was significantly higher in divisions 3-6 of OT-I T cells in WT mice compared to those from *Batf3*^-/-^ mice (Figure 7K-M) in line with the most significant difference seen within the model. These data in conjunction with the agent-based model indicated that cDC1 are important to promote entry of CD8^+^ T cells into pro-survival blastogenesis after T cell priming, and that the loss of cDC1 increased the rate of T cell death and resulted in a defect in the expression of nutrient and IL-2 receptors that support blastogenesis (*54*-*56*).

## DISCUSSION

The importance of WAT and the FALCs located therein to the immune response against infection is increasingly appreciated (*4, 13, 57*). In the present study, we have shown that after CPS immunization the MS in the omentum are a major site of antigen drainage as well as the site of T cell priming and expansion. One of the primary functions of the omentum is the collection of antigens leaving the peritoneum via convective flow or trafficking by innate immune cells through stomata in the mesothelial sheet into the vascularized MS (*11, 12, 14*-*19*). The structure and placement of these lymphoid aggregates at a serosal surface facilitates interactions between innate and adaptive immune cells trafficking to the MS from both the blood and the peritoneum. The coordinated response within the MS to CPS immunization results in a marked increase and change in cellularity that illustrates the dynamic nature of the MS within the omentum. Utilization of Cre-secreting CPS parasites revealed that infected peritoneal macrophages migrate from the peritoneum into the MS rather than to the draining LN. While the factors that initiate these changes in the MS are not well understood, recent work has implicated stromal cell production of chemokines in the recruitment of innate lymphoid cells to the MS (*15, 19*). However, the process of mobilization of peritoneal macrophages induced by *T. gondii* is known to require IFN-*γ* produced by type I ILCs (*58*), which can be activated by IL-12 produced by cDC1 (*40, 59*). Indeed, it is shown here that changes in the cellularity of the omentum are also dependent on the early immune response initiated by cDC1. Counterintuitively, infection of Ai6/*Batf3*^-/-^ mice revealed that the accumulation of infected macrophages in the MS was independent of cDC1. This migration of infected macrophages into the MS could result from parasite effector proteins that enhance the motility and ability of infected macrophages and DC to invade tissues (*60*-*63*).

In uninflamed tissue the reports that FALCs lack organized T and B cell regions, follicular dendritic cells (FDCs), and a central reticular network have led to the idea that MS are not SLOs (*12*). Previous work has shown that adaptive immune responses thought to occur exclusively in SLOs can be initiated in MS of the omentum to drive T cell-dependent B cell responses to vaccination (*12, 15*) and T cell responses to male histocompatibility antigen (*16*), but T cell responses to microbial challenge have remained unexplored. Here the use of multiple approaches establishes that in response to i.p. microbial challenge CD8^+^ T cells are not only primed but also expand in the MS of the omentum. The current paradigm for many pathogens is that *infected* migratory cDCs deliver antigen to SLOs and that their position in SLOs is critical to initiate T cell activation (*25*). Similarly, high-resolution imaging of the MS utilized here showed that infected macrophages migrate into distinct T and B cell zones, and that priming occurred in the T cell zones associated with cDC1. Thus, the MS and perhaps FALCs in other WAT should be considered as a relevant site for the initiation of T and B cell mediated immunity.

The role for cDC1 in adipose tissue is not well understood. Previous work has shown a role for cDC1 in the WAT in controlling whole body homeostasis (*26*), while priming of CD8^+^ T cells in the omentum was attributed to CD11b^+^ cDC2s (*16*). Imaging of the MS revealed that cDC1 were enriched in the T cell zones similar to the organization of cDC1 in LNs (*24*), where they were closely associated with clusters of activated OT-I T cells. In immunized *Batf3*^-/-^ mice, the loss of the parasite-specific CD8^+^ T cell response indicated that cDC1 in the MS played a critical role in generating protective cellular immunity. The major unanticipated phenotype is that cDC1 are not required for priming of CD8^+^ T cells (*36*), but rather were essential for the clustering of activated T cells and subsequently the full expansion of pathogen-specific responses. These data indicated that open questions remain as to the role of cDC1 in generating the CD8^+^ T cell response to CPS immunization. One possible explanation is that the CD8^+^ T cell response generated by CPS immunization is dependent on CD4^+^ T cell help (*31*) similar to other models (*21, 64*). cDC1 can serve as a platform for the interaction between CD8^+^ and CD4^+^ T cells to facilitate T cell help (*21, 22*). These studies demonstrate that in addition to their role in maintaining homeostasis, cDC1 in WAT are capable of functioning in the same capacity as their counterparts in SLOs to drive robust CD8^+^ T cell responses.

The requirements for T cell activation and clonal expansion have been well studied *in vitro*, but the redundancy of extrinsic signals makes it difficult to translate them to *in vivo* T cell responses. To better understand this process, we utilized the agent-based STORE model of OT-I T cell activation and expansion in the omentum. Modeling of the adaptive immune response has been a useful tool to understand T cell differentiation pathways *in vivo* (*53, 65*-*67*), as well as the factors that determine the division destiny of lymphocytes *in vitro* (*68*-*71*). However, these deterministic models do not account for randomness in biological samples and are poorly suited for the analysis of small numbers of cells or to capture rare events such as initial APC-T cell interactions (*72*). Importantly, this stochastic model required only a small number of parameters to recapitulate the early *in vivo* response of OT-I T cells in WT and *Batf3*^-/-^ recipients and did not appear sensitive to the initial number of naïve OT-I T cells. The model correctly predicted an increased rate of cell death of OT-I T cells in their early divisions in the *Batf3*^-/-^ mice, which was also characterized by decreased expression of nutrient uptake receptors required for survival and blastogenesis of T cells (*54*-*56*). While the studies presented here establish the role of cDC1 in the co-ordination of the innate and adaptive immune response in atypical secondary lymphoid structures in WAT, they also highlight a critical window following activation in which CD8^+^ T cell:cDC1 interactions are crucial for the generation of protective CD8^+^ T cell responses and novel questions about additional contributions of cDC1 outside of antigen cross-presentation and IL-12 production that have important implications for T cell responses. The development of an accurate stochastic model of early T cell responses is a powerful tool that provides a clearer picture of the response at timescales far faster than experiments are able to probe, which will allow for the directed design of vaccination strategies that elicit a protective cellular immune response.

## ACKNOWLEDGEMENTS

We would like to acknowledge the contributions of the Andrea Stout, PhD and the Cell and Developmental Biology Microscopy Core at the Perelman School of Medicine, University of Pennsylvania. We would also like to acknowledge the contributions of the Penn Vet Imaging Core at the University of Pennsylvania School of Veterinary Medicine. This work was supported by grants from the National Institutes of Health: NIAID AI126899-01 RO1/UO1, NIAID 1R21AI126042-01, NIAID R01AI125563.

## AUTHOR CONTRIBUTIONS

Conceptualization, D.A.C., T.A.A., A.T.P., and C.A.H.; Methodology, D.A.C. and T.A.A.; Software, T.A.A., T.E.S., and M.A.; Formal Analysis, D.A.C., T.A.A. and T.E.S.; Investigation, D.A.C., T.A.A., L.A.S., M.A., J.P., and J.T.C; Resources, D.J.T. and K.M.M; Writing – Original Draft, D.A.C. and C.A.H.; Writing – Review & Editing, D.A.C., A.T.P., and C.A.H.; Visualization, D.A.C. and G.R.; Supervision, D.A.C. and C.A.H.; Project Administration, D.A.C. and C.A.H.; Funding Acquisition, R.M.K and C.A.H.

## SUPPLEMENTAL INFORMATION

Figure S1. Visualizing recruitment of LysM^+^ myeloid cells to the MS

Figure S2. Immunization-induced recruitment of monocytes to the peritoneum requires cDC1 and IL-12p40

Figure S3. Mitochrondrial mass and glycolytic uptake measurements of OT-I T cells in WT and *Batf3*^-/-^ mice

Figure S4. K-function statistical analysis of T cell clustering.

Figure S5. K-function statistical analysis of clustering.

Figure S6. Cross K-function statistical analysis of T cell clustering.

Figure S7. Cross K-function statistical analysis of clustering. Statistical Tests of Clustering

Supplemental Movie 1

## MATERIALS AND METHODS

### Ethics statement

All procedures involving mice were reviewed and approved by the Institutional Animal Care and Use Committee of the University of Pennsylvania (Animal Welfare Assurance Reference Number #A3079-01) and were in accordance with the guidelines set forth in the Guide for the Care and Use of Laboratory Animals of the National Institute of Health.

### Mice

*Batf3*^-/-^, *Il12b*^-/-^, Ai6, Nur77^GFP^, OT-I, and LysM-Cre mice were obtained from Jackson Laboratories. C57BL/6, Rag2^-/-^, and Rag2^-/-^*γ*c^-/-^ mice were obtained from Taconic Farms. CD45.1 mice were obtained from Charles River Laboratories. *Snx22*^GFP/GFP^ mice were on a 129S1 background and were obtained from Dr. Kenneth M. Murphy at Washington University in St. Louis. *Snx22*^GFP/+^ were an F1 cross with C57BL/6 mice. For IL-12p70 add back experiments, C57BL/6 and *Batf3*^-/-^ mice were treated with an intraperitoneal (i.p.) injection of either PBS (Corning: 21-0311-CM) or 200 ng of recombinant murine IL-12p70 (Peprotech: 210-12) immediately after immunization and once per day for the next two days after immunization. For 2-NBDG uptake assays, mice were injected i.p. with 100 μL of 0.5 mM 2-NBDG (ThermoFisher Scientific: N13195) 30 min prior to euthanization. All mice were kept in specific-pathogen-free conditions at the School of Veterinary Medicine at the University of Pennsylvania.

### Immunizations

All experiments were performed using *cpsII* parasites, *cpsII*- OVA parasites, or *cpsII*-Cre-OVA-mCherry parasites. *CpsII*-OVA parasites have been previously described (*73*) and were derived from the RHΔ*cpsII* clone, which was provided as a generous gift by Dr. David Bzik (*28*). *CpsII*-Cre-OVA-mCherry parasites were derived from the *cpsII*-OVA clone using the previously described methods (*37, 74*) with the exception that parasites were selected using zeomycin as previously described (*75*). Parasites were cultured and maintained by serial passage on human foreskin fibroblast cells in the presence of parasite culture media [71.7% DMEM (Corning: 10-017-CM), 17.9% Medium 199 (Gibco: 11150-059), 9.9% Fetal Bovine Serum (FBS)(Atlanta Biologics: S11150H), 0.45% Penicillin and Streptomycin (Gibco: 15140-122)(final concentration of 0.05 units/ml Penicillin and 50 µg/ml Streptomycin), 0.04% Gentamycin (Gibco: 15750-060)(final concentration of 0.02 mg/ml Gentamycin)], which was supplemented with uracil (Sigma-Aldrich: U1128) (final concentration of 0.2 mM uracil). For infections, parasites were harvested and serially passed through 18 and 26 gauge needles (BD: 305196, 305115) before filtration with a 5 µM filter (PALL Acrodisc: 4650). Parasites were washed extensively with PBS and mice were injected i.p. with parasites suspended in PBS.

### T cell transfers and tissue harvesting

For T cell transfers OT-I mice were crossed with CD45.1/Nur77^GFP^ mice. To isolate OT-I CD8^+^ T cells, lymph nodes and spleen were harvested and leukocytes from the spleen and draining lymph nodes were obtained by processing spleens and lymph nodes over a 70 µm filter (Fisher Scientific: 22-363-548) and washing them in complete RPMI [90% RMPI 1640 (Corning: 10-040-CM), 10% FBS, 1% penicillin-streptomycin, 1 mM sodium pyruvate (Corning: 25-000-Cl), 1% nonessential amino acids (Gibco: 11140-050), and 0.1% *β*-mercaptoethanol (Gibco: 21985-023)]. Red blood cells were then lysed by incubating for 5 minutes at room temperature in 5 ml of lysis buffer [0.864% ammonium chloride (Sigma-Aldrich: A0171) diluted in sterile de-ionized H2O)], followed by washing with complete RPMI. OT-I CD8^+^ T cells were then purified by magnetic activated cell sorting (MACS) using the CD8a+ T Cell Isolation Kit (Miltenyi Biotec: 130-104-075). For cell division or microscopy studies, purified OT-I T cells were then fluorescently labeled using the CellTrace Violet labeling kit (ThermoFisher Scientific: C34557). OT-I T cells were then transferred by i.p. injection into recipient mice. Peritoneal exudate cells were obtained by peritoneal lavage with 8 mL of ice cold PBS. Omentum was isolated, incubated in 0.4 U/mL of LiberaseTL (Roche: 5401020001) for one hour at 37°C, passed through an 18G needle, and processed over a 70 μm filter. Leukocytes from the spleen and draining lymph nodes were obtained by processing spleens and lymph nodes, washing them in complete media, and lysing red blood cells (see above). Cells were then resuspended in complete RPMI.

### Flow cytometry

Cells were washed with FACS buffer [1× PBS, 0.2% bovine serum antigen (Gemini: 700-100P), 1 mM EDTA (Gibco: 15575-038)] and incubated in Fc block [99.5% FACS Buffer, 0.5% normal rat IgG (Invitrogen: 10700), 1 µg/ml 2.4G2 (BioXCell: BE0307)] at 4°C for 10 min prior to staining. If cells were stained for cell death using LIVE/DEAD staining, LIVE/DEAD Fixable Aqua Dead Cell marker (Invitrogen: L34957) was included during incubation with Fc block. Tetramer-specific CD8^+^ and CD4^+^ T cells were measured by staining in 50 μL FACS buffer containing MHCI and MHCII tetramers specific for endogenous *T. gondii* antigens for 1 hour at room temperature. Cells were surface stained in 50 μL at 4°C for 15-20 min and washed in FACS buffer prior to acquisition. For intracellular cytokine staining, 10^6^ cells from the peritoneal exudate or omentum were plated in a 96-well plate and incubated with 1X Brefeldin A (Sigma-Aldrich: B7651) at 37°C for 3 hr. For intracellular cytokine and transcription factor staining, cells were rinsed with FACS buffer and surface stained as described above, fixed using the eBioscience Foxp3 Transcription Factor Fixation/Permeabilization Concentrate and Diluent (ThermoFisher Scientific: 00-8222) for 30 min at 4°C, and then washed with FACS buffer. Cells were then stained for intracellular cytokines and transcription factors in 50 μL 1X eBioscience Permeabilization Buffer (ThermoFisher Scientific: 00-8333-56) at 4°C for at least 1 hr. Cells were then washed in FACS buffer prior to acquisition. The APC Annexin V Kit (Biolegend: 640920) was used to measure cell death via Annexin V and propidium iodide (PI). Cells were surface stained and washed with FACS buffer then resuspended in 100 μL Annexin V binding buffer (Biolegend: 79998). Cells were then incubated with 5 μL APC Annexin V (Biolegend: 640920) and 10 μL PI solution (Biolegend: 79997) for 15 min at room temperature in the dark. Prior to acquisition, 400 μL of Annexin V binding buffer was added to each sample. For mitochondrial staining 2 × 10^6^ cells were suspended in 100 μL HBSS (Gibco: 14175-079) supplemented with 0.5 mM MgCl2 (Sigma-Aldrich: M4880). Mitochondrial stains were then added at the following concentrations: 50 nM MitoTracker Green FM (Invitrogen: M7514), 25 nM MitoTracker Deep Red FM (Invitrogen: M22426) and 5 mM MitoSOX Red (Invitrogen: M36008). Samples were incubated at 37°C for 30 min, rinsed with FACS buffer, and surface stained as above prior to acquisition. The following antibodies were used for staining: B220: BV510, Biolegend: 103248, clone: RA3-6B2; B220: ef450, eBioscience: 48-0452-82, clone: RA3-6B2; B220: PerCp-Cy5.5, Biolegend: 103236, clone: RA3-6B2; CCR2: AF647, Biolegend: 150604, clone: SA203G11; CD102: BUV395, BD Biosciences: 740227, clone: 3C4 (mIC2/4); CD102: ef450, eBioscience: 48-1021-82, clone: 3C4 (mIC2/4); CD102: FITC, Biolegend: 105606, clone: 3C4 (mIC2/4); CD115: BV605, Biolegend: 135517, clone: AF398; CD11a: PerCp-Cy5.5, Biolegend: 101124, clone: M17/4; CD11b: APC-ef780, eBioscience: 47-0112-82, clone: M1/70; CD11b: BV605, Biolegend: 101247, clone: M1/70; CD11b: BV650, Biolegend: 101259, clone: M1/70; CD11c: APC-Fire750, Biolegend: 117352, clone: N418; CD11c: APC-R700, BD Biosciences: 565872, clone: N418; CD11c: BUV737, BD Biosciences: 612797, clone: HL3; CD172a: APC, Biolegend: 144014, clone: P84; CD172a: APC-Cy7, Biolegend: 144018, clone: P84; CD19: BV605, Biolegend: 115539, clone: 6D5; CD19: ef450, eBioscience: 48-0193-82, clone: eBio1D3; CD19: PerCp-Cy5.5, Biolegend: 152406, clone: ID3/CD19; CD200R: APC, eBioscience: 17-5201-82, clone: OX110; CD25: APC, eBioscience: 17-0251-82, clone: PC61.5; CD25: PE-Cy7, eBioscience: 25-0251-82, clone: PC61.5; CD3: APC-ef780, Invitrogen: 47-0032-82, clone: 17A2; CD3: BV605, Biolegend: 100237, clone: 17A2; CD3: BV750, Biolegend: 100249, clone: 17A2; CD3: BV785, Biolegend: 100232, clone: 17A2; CD3: PacBlue, Biolegend: 100214, clone: 17A2; CD3e: PE-cf594, BD Biosciences: 562286, clone: 145-2C11; CD3e: PerCp-Cy5.5, Invitrogen: 45-0031-82, clone: 145-2C11; CD335: BV650, Biolegend: 137635, clone: 29A1.4; CD335: PE-Dazzle594, Biolegend: 137630, clone: 29A1.4; CD4: AF700, Biolegend: 100536, clone: RM4-5; CD4: BV650, Biolegend: 100555, clone: RM4-5; CD4: BV711, Biolegend: 100447, clone: GK1.5; CD4: BV785, Biolegend: 100552, clone: RM4-5; CD44:BV605, BD Biosciences: 563058, clone: IM7; CD44: BV785, Biolegend: 103059, clone: IM7; CD45: BUV805, BD Biosciences: 748370, clone: 30-F11; CD45.1: APC-ef780, Invitrogen: 47-0453-82, clone: A20; CD45.1: BV711, Biolegend: 110739, clone: A20; CD45.1: ef450, Invitrogen: 48-0453-82, clone: A20; CD45.1: PE, BD Biosciences: 553776, clone: A20; CD45.1: PE-cf594, BD Biosciences: 562452, clone: A20; CD45.1: PerCp-Cy5.5, Invitrogen: 45-0453-82, clone: A20; CD45.2: APC-ef780, Invitrogen: 47-0454-82, clone: 104; CD45.2: BV711, Biolegend: 109847, clone: 104; CD49b: FITC, BD Biosciences: 553857, clone: DX5; CD5: ef450, eBioscience: 48-0051-82, clone: 53-7.3; CD5: PerCp-Cy5.5, eBioscience: 45-0051-82, clone: 53-7.3; CD62L: APC-ef780, eBioscience: 47-0621-82, clone: MEL-14; CD62L: BV711, Biolegend: 104445, clone: MEL-14; CD64: PE-Cy7, Biolegend: 139314, clone: X54-5/7.1; CD69: BV711, Biolegend: 104537, clone: H1.2F3; CD69: ef450, Invitrogen: 48-0691-82, clone: H1.2F3; CD69: PE-Cy7, BD Biosciences: 552879, clone: H1.2F3; CD69: PerCp-Cy5.5, eBioscience: 45-0691-82, clone: H1.2F3; CD71: PE, Biolegend: 113807, clone: R17217; CD8a: BV605, Biolegend: 100744, clone: 53-6.7; CD8a: BV650, Biolegend: 100742, clone: 53-6.7; CD8a: BV711, Biolegend: 100747, clone: 53-6.7; CD8b: APC-ef780, Invitrogen: 47-0083-82, clone: ebioH35-17.2; CD8b: FITC, Biolegend: 126606, clone: YTS156.7.7; CD90.2: AF700, Biolegend: 105320, clone: 30-H12; CD98: PE-Cy7, Biolegend: 128214, clone: RL388; ckit: APC-ef780, Invitrogen: 47-1172-80, clone: ACK2; CXCR3: BV650, Biolegend: 126531, clone: CXCR3-173; CXCR3: PE-Cy7, Biolegend: 126516, clone: CXCR3-173; EOMES: PE, Invitrogen: 12-4875-80, clone: Dan11mag; F4/80: ef450, Invitrogen: 48-4801-80, clone: BM8; FcεR1a: APC, Invitrogen: 17-5898-82; clone: 1-Mar; Glut1: AF647, AbCam: ab195020, clone: EPR3915; I-A/I-E: AF700, Biolegend: 107622, clone: M5/144.15.2; I-A/I-E: BV711, Biolegend: 107643, clone: M5/144.15.2; IFN-*γ*: PE-Cy7, Invitrogen: 25-7311-82, clone: XMG1.2; KLRG1: BV711, Biolegend: 138427, clone: 2F1/KLRG1; KLRG1: FITC, eBioscience: 11-5893-82, clone: 2F1; LFA1: PerCp-Cy5.5, Biolegend: 141008, clone: H155-78; Ly6C: BV570, Biolegend: 128030, clone: HK1.4; Ly6C: BV785, Biolegend: 128041, clone: HK1.4; Ly6G: BUV563, BD Biosciences: 612921, clone: 1A8; Ly6G: PE, Biolegend: 127608, clone: 1A8; NK1.1: BV711, Biolegend: 108745, clone: PK136; NK1.1: PerCp-Cy5.5, Biolegend: 108728, clone: PK136; SiglecF: PE, BD Biosciences: 552126, clone: E50-2440; SiglecH: BV421, BD Biosciences: 566581, clone: 440C; TCR V*α*2: PE, Biolegend: 127808, clone: B20.1; TCR V*α*2: Superbright780, Invitrogen: 78-5812-82, clone: B20.1; TCR V*β*5.1,5.2: APC, Biolegend: 139506, clone: MR9-4; TCR*α*: PerCp-Cy5.5, Biolegend: 109228, clone: H57-597; Tetramer MHCI: PE, NIH Tetramer Core, peptide: SVLAFRRL; Tetramer MHCII: PE, NIH Tetramer Core, peptide: AVEIHRPVPGTAPPS; XCR1: BV650, Biolegend: 148220, clone: ZET; XCR1: BV785, Biolegend: 148225, clone: ZET. Samples were run on a LSR Fortessa (BD), LSRII (BD), or FACSymphony A5 (BD) and analyzed using FlowJo Software (TreeStar).

### Immunofluorescence imaging

Whole omenta were harvested from mice and fixed in 1% PFA overnight at 4°C. After rinsing, tissue was blocked using 10% BSA, 0.5% normal rat serum (Invitrogen), and 1 µg/ml 2.4G2 (BD) in PBS for 1 hr at room temperature. For immunofluorescence staining, omenta were incubated in PBS containing primary antibodies at 4°C for 3 days and subsequently rinsed with PBS overnight. Antibodies included: B220: AF647, BD Biosciences: 557683, clone: RA3-6B2; B220: BV510, Biolegend: 103248, clone: RA3-6B2; CD11b: AF700, eBiosciences: 56-0112-82, clone: M1/70; CD3: AF532, Invitrogen: 58-0032-82, clone: 17A2; CD4: AF532, Invitrogen: 58-0042-82, clone: RM4-5; CD8b: ef450, Invitrogen: 48-0082-82, clone: eBioH35-17.2; ER-TR7: AF647, Santa Cruz Biotechnology: sc-73355 AF647, clone: ER-TR7. Tissues were then mounted on slides using Prolong Diamond (Invitrogen: P36970) or Prolong Glass mounting media (Invitrogen: P36984) and imaged using a Leica TCS SP8 STED microscope and acquired using LAS X (Leica Microsystems). Images were processed and analyzed using LAS X and Imaris (Oxford Instruments).

### Cluster analysis

Clustering was analyzed using K function statistics as described in full in Appendix C. Briefly, to determine clustering of Nur77^GFP+^ OT-I T cells, the average number of Nur77^GFP+^ cells within a specified distance, *k*, of each Nur77^GFP+^ cell was compared with those averages calculated for 999 randomly permuted patterns of the full population of both Nur77^GFP+^ and Nur77^GFP-^ cells. The subpopulation of Nur77^GFP+^ cells was then deemed to be significantly clustered at scale *k* if the fraction of patterns with averages as large as the observed pattern was not greater than 0.05. To determine whether Nur77^GFP+^ cells were clustered around a specific APC population, the target population is deemed to be significantly clustered around the reference population at scale *k* if the fraction of randomly permuted populations with an average number of target cells with *k* as large as the observed population is not larger than 0.05. Similarly, the target population is deemed to be significantly dispersed away from the reference population at scale *k* if the fraction of populations with averages as small as the observed population is not larger than 0.05.

### STochastic Omentum REsponse (STORE) model

The STORE model is a dynamic stochastic mathematical model used to simulate the early OT-I T cell response to immunization in the omentum. Briefly, the model is based off experimental data measuring the kinetics of the OT-I T cell response in the omentum through the first 8 days of immunization. Simulations of the model track the interactions of transferred OT-I T cells with APC populations and the subsequent T cell divisions that drive the expansion of the OT-I T cell population. The behavior of every cell within the system is tracked individually throughout the simulation and is determined by probability distributions estimated from the literature or taken as a separate parameter. The goal of the model is to recapitulate the experimental data for OT-I T cells with the tabulated parameters defining the differences between the responses of OT-I T cells in the WT and *Batf3*^-/-^ mice.

### Statistical analysis

Statistical analysis was performed using PRISM software (Graphpad Software). Significance was calculated using an unpaired two-tailed student’s t-test.

