## Supplemental Information for "cDC1 Coordinate Innate and Adaptive Responses in the Omentum required for T cell Priming and Memory"

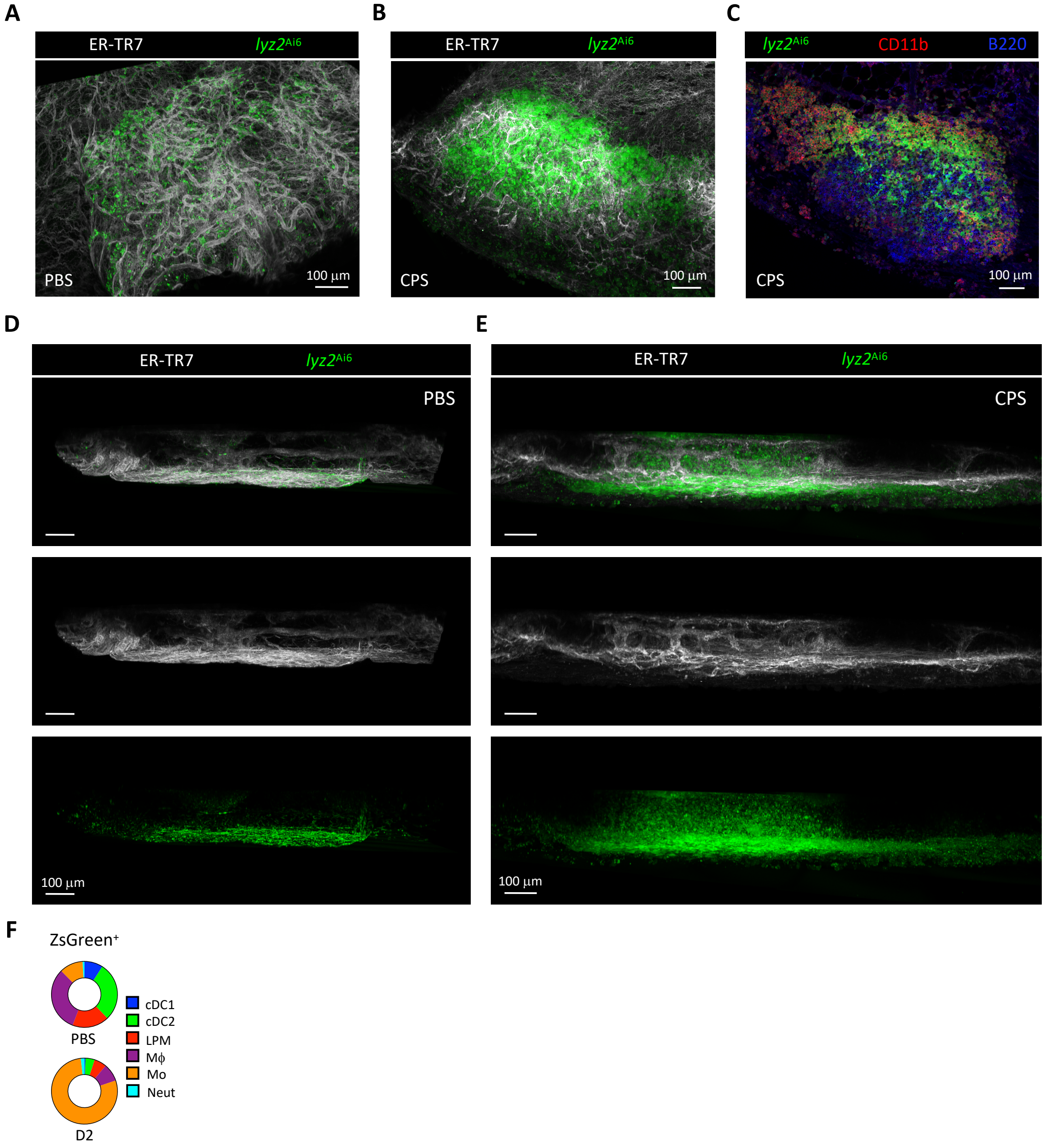


**Figure S1. Visualizing recruitment of LysM$^+$ myeloid cells to the MS**

(A-E) Immunofluorescence staining of omentum MS from *lyz2*$^{Ai6}$ mice. (A and B) 3D rendering of MS showing enrichment of ER-TR7$^+$ reticular cells and reticular fibers in the MS in PBS treated and CPS-immunized ($10^5$ parasites) mice at 2 dpi. Immunization results in increased recruitment of ZsGreen$^+$ myeloid cells. ER-TR7 (gray), ZsGreen (green).

(C) Z-slice of MS showing recruitment of CD11b$^+$ZsGreen$^+$ myeloid cells to the MS and forming a mantle around the lymphocyte-rich MS after CPS immunization (2 dpi). ZsGreen (green), CD11b (red), B220 (blue).

(D and E) 3D renderings of MS on XZ plane showing ZsGreen$^+$ myeloid cells remain on the surface of the omentum after PBS treatment while infiltrating into the MS after CPS immunization (2 dpi). ER-TR7 (gray), ZsGreen (green).

(F) Donut charts represent the ZsGreen$^+$ cellular distribution from PBS treated and CPS-immunized ($10^5$ parasites, 2 dpi) *lyz2*$^{Ai6}$ mice. cDC1 (Lin$^-$CD64$^-$CD11c$^+$MHCII$^+$XCR1$^+$CD172a$^-$), cDC2 (Lin$^-$CD64$^-$CD11c$^+$MHCII$^+$XCR1$^-$CD172a$^+$), LPM (Lin$^-$CD64$^+$CD11b$^+$CD102$^+$), Mϕ (Lin$^-$CD64$^+$CD11b$^+$CD102$^-$Ly6G$^-$Ly6C$^-$), Mo (Lin$^-$CD64$^+$CD11b$^+$CD102$^-$Ly6G$^-$Ly6C$^+$), Neut (Lin$^-$CD64$^-$CD11c$^-$CD11b$^+$Ly6G$^+$Ly6C$^{int}$). Lin (CD3, CD19, B220, NK1.1).

Scale bars in all images are 100 µm.

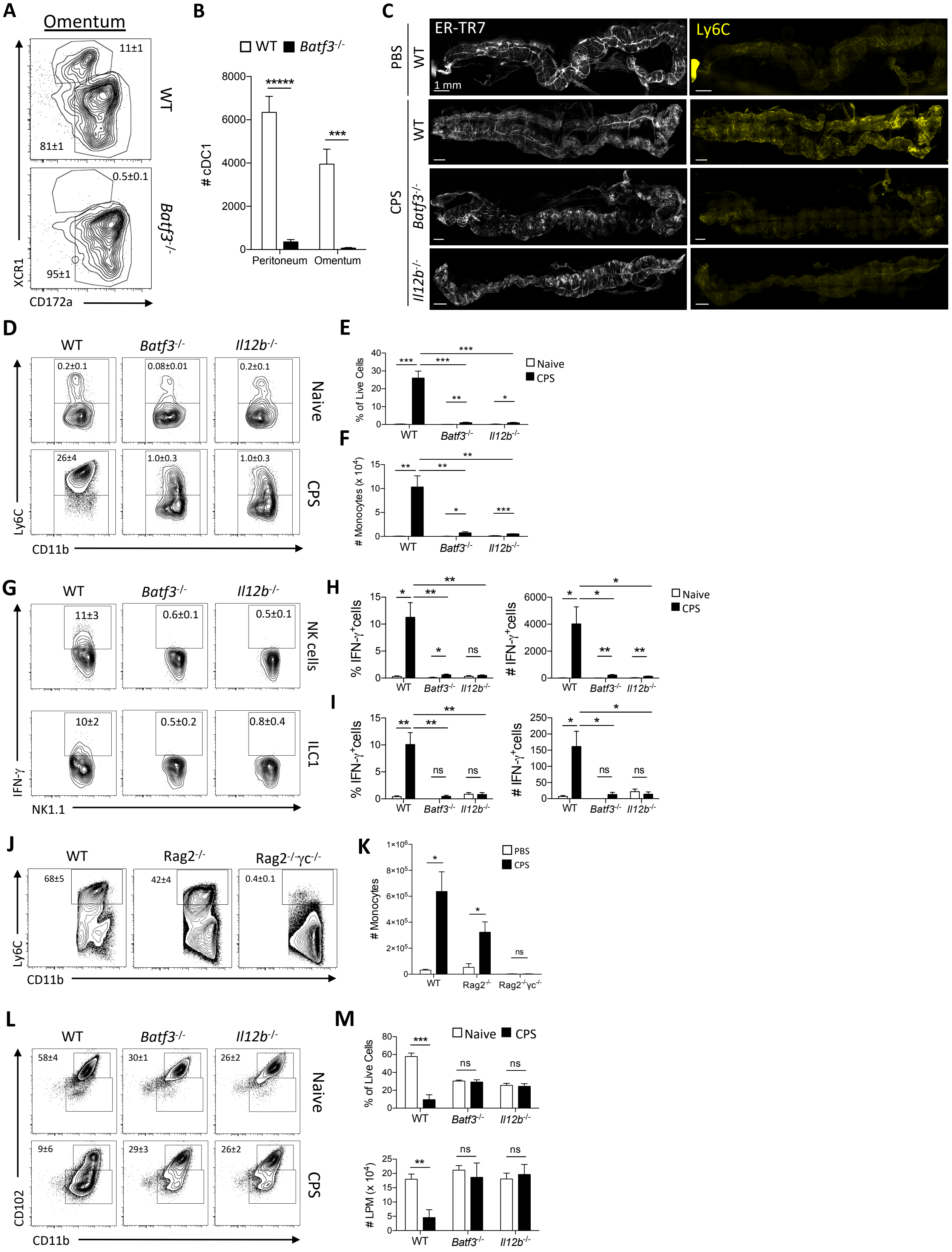

A
Omentum
11±1
WT
81±1
0.5±0.1
Batf3-/-
95±1
XCR1
CD172a
B
WT Batf3-/-
### cDC1
Peritoneum
Omentum
C
ER-TR7
Ly6C
PBS
WT
1 mm
CPS
Batf3-/-
Il12b-/-
D
Naive
CPS
Ly6C
CD11b
E
% of Live Cells
Naive
CPS
F
### Monocytes (x 10^4)
G
NK cells
ILC1
IFN-γ
NK1.1
H
% IFN-γ+cells
### IFN-γ+cells
I
J
Rag2-/-
Rag2-/-γc-/-
K
PBS
CPS
### Monocytes
L
CD102
CD11b
M
### LPM (x 10^4)

**Figure S2. Immunization-induced recruitment of monocytes to the peritoneum requires cDC1 and IL-12p40**

(A-B) Quantification of cDC1 fraction and numbers in the omentum of WT and *Batf3*$^{-/-}$ mice immunized with $10^5$ CPS parasites at 2 dpi. (A) Flow cytometric analysis of cDC1 in the omentum (Lin$^-$CD64$^-$CD11c$^+$MHCII$^+$XCR1$^+$CD172a$^-$). Lin (CD3, CD19, B220, NK1.1)
(B) Quantification of the number of cDC1 in the peritoneum and omentum. Data (n = 4-5) represent 1 out of 5 independent experiments.
(C) Immunofluorescence images of omenta from WT, *Batf3*$^{-/-}$, and *Il12b*$^{-/-}$ mice showing recruitment of Ly6C$^+$ cells to the omentum after CPS immunization (2 dpi) is dependent upon cDC1 and IL-12p40. ER-TR7 (gray), Ly6C (yellow). Scale bars are all 1 mm.
(D-F) Quantification of recruitment of monocytes to the peritoneum of WT, *Batf3*$^{-/-}$, and *Il12b*$^{-/-}$ mice after immunization with $10^5$ CPS parasites (2 dpi). (D) Flow cytometry analysis of monocytes (CD64$^+$CD11b$^+$CD102$^-$Ly6C$^{HI}$) in the peritoneum.
(E and F) Quantification of monocytes as a fraction of live cells in the peritoneum and the total number of monocytes in the peritoneum. Data (n = 4-5) represent 1 experiment out of 2-3 independent experiments.
(G-I) Quantification of IFN-γ production by Type I ILCs in the peritoneum of WT, *Batf3*$^{-/-}$, and *Il12b*$^{-/-}$ mice after immunization with $10^5$ CPS parasites (1 dpi). (G) Flow cytometry analysis of NK cells (Lin$^-$NK1.1$^+$NKp46$^+$EOMES$^+$CD200R$^-$) and ILC1s (Lin$^-$NK1.1$^+$NKp46$^+$EOMES$^-$CD200R$^+$) producing IFN- γ.
(H and I) Quantification comparing the fraction and total number of (H) NK cells and (I) ILC1s producing IFN-γ between naive and CPS immunized mice at 1 dpi. Lin (CD3, CD5, CD19, B220, F4/80). Data (n = 4-5) represent 1 experiment out of 2-3 independent experiments.
(J and K) Quantification of monocyte recruitment to the peritoneum of WT, Rag2$^{-/-}$, and Rag2$^{-/-}$γc$^{-/-}$ mice after PBS treatment or immunization with $10^5$ CPS parasites (2 dpi). (J) Flow cytometric analysis of monocytes (Lin$^-$CD11b$^+$Ly6G$^-$CD90.2$^-$Ly6C$^{HI}$) in the peritoneum. Lin (CD3, CD5, B220, CD19).
(K) Quantification of total number monocytes in the peritoneum. Data (n = 3-5) represent 1 experiment.
(L and M) Quantification of the loss of LPM from the peritoneum induced by immunization with $10^5$ CPS parasites (2 dpi) in WT, *Batf3*$^{-/-}$, and *Il12b*$^{-/-}$ mice. (L) Flow cytometric analysis of LPM (CD64$^+$CD11b$^+$CD102$^+$) in the peritoneum.
(M) Quantification of the LPM in the peritoneum as a fraction of total live cells and total number of LPM in naïve and CPS-immunized mice. Data (n = 3-5) represent 1 experiment out of 2-3 independent experiments.
All representative plots indicate mean ± SD. All statistical comparisons were unpaired Student's t test. ns, not significant; *$p < 0.05$; **$p < 0.01$; ***$p < 0.001$; *****$p < 0.00001$.

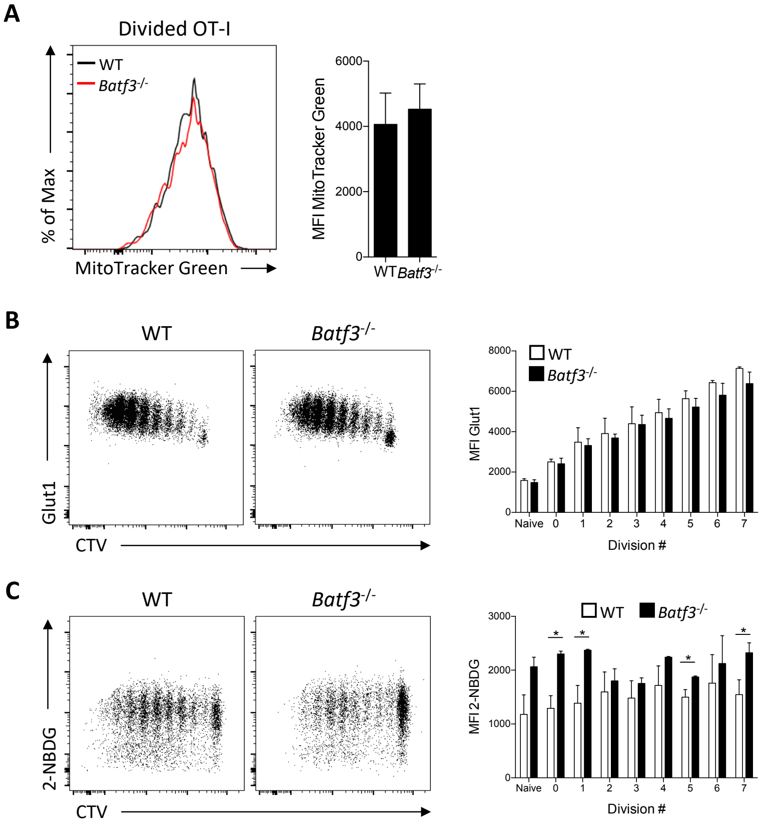


**Figure S3. Mitochrondrial mass and glycolytic uptake measurements of OT-I T cells in WT and *Batf3*$^{-/-}$ mice**
(A-C) Intraperitoneal transfer of 5 x $10^5$ CTV-labeled OT-I/Nur77$^{GFP}$ T cells into WT and *Batf3*$^{-/-}$ mice immunized with 2 x $10^5$ CPS-OVA parasites 2 hr later. OT-I T cells in the omentum at 3 dpi were evaluated by flow cytometry. (A) Representative plot of the intensity of MitoTracker Green, and quantification of the MFI of MitoTracker Green in OT-I T cells that had undergone one or more divisions. Data (n = 4-5) represent 1 experiment out of 2 independent experiments.
(B) Representative plot of the expression of Glut1, and quantification of the MFI of Glut1 on OT-I T cells. Data (n = 3-5) represent 1 experiment out of 2 independent experiments.
(C) Representative plot of the uptake of 2-NBDG, and quantification of the MFI of 2-NBDG in OT-I T cells. Data (n = 3) represent 1 experiment out of 2 independent experiments.
All representative plots indicate mean ± SD. All statistical comparisons were unpaired Student's t test. *$p < 0.05$.

#### Statistical Test of Clustering

##### Tests using K-functions

The following test procedure is best illustrated by an example from the text. The results in Figure 3G of the text summarize clustering tests for activated cells, Nur77$^{GFP+}$, relative to non-activated cells, Nur77$^{GFP-}$, for nine different milky spots in the omentum of two mice. One of these test cases is illustrated in Figure S4A.

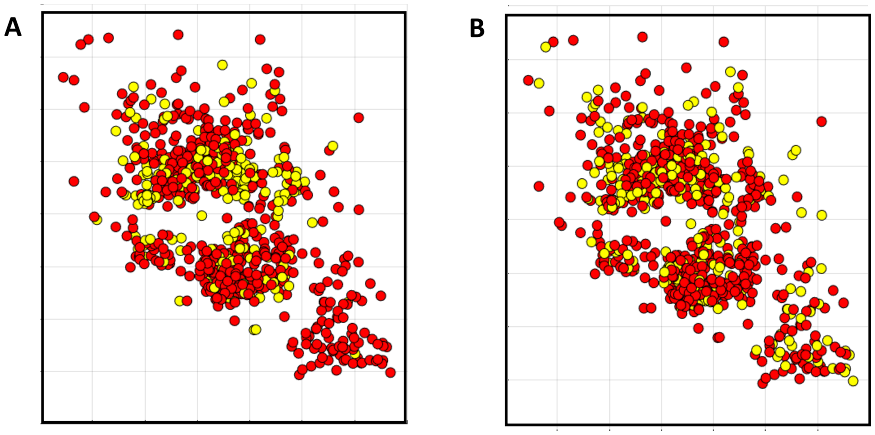


**Figure S4. K-function statistical analysis of T cell clustering.**
(A-B) Intraperitoneal transfer of 5 x $10^5$ CTV-labeled OT-I/Nur77$^{GFP}$ T cells into WT mice immunized with 2 x $10^5$ CPS-OVA parasites 2 hr later. Immunofluorescent images were analyzed to determine the coordinates of OT-I T cells. (A) 2-dimensional (2D) model representation of coordinates of Nur77$^{GFP+}$ (yellow) and Nur77$^{GFP-}$ (red) OT-I T cells in a MS.
(B) Representative randomized distribution of Nur77$^{GFP+}$ and Nur77$^{GFP-}$ OT-I T cells from (A).

This figure shows the observed pattern of $n_1 = 239$ Nur77$^{GFP+}$ cells (yellow) together with $n_2 = 613$ Nur77$^{GFP+}$ cells (red) in the same milky spot. This Matlab coordinate plot is a 2D projection of a 3D sample. Because this sample slice has a thickness of only 50 $\mu m$ (relative to the box width of 630 $\mu m$ in Figure S4A) such 2D projections convey almost all relevant spatial information. The question of interest is whether the subpopulation of yellow (activated) cells is significantly more clustered that a typical random subpopulation of $n_1$ cells within the total population of $n = n_1 + n_2 = 852$ cells. One such random subpopulation of $n_1$ yellow cells is shown in Figure S4B. A visual comparison of the two suggests that the observed subpopulation on the left is indeed more clustered, and in particular contains almost no cells in the lower right corner, unlike the random subpopulation.

To test this more formally, we start by letting $P^0 = \{x_i : i = 1,..,n\}$ denote the set of observed points (with 3D coordinate locations) in Figure S4A, where the first $n_1$ points are taken to represent the target subset of "activated" points, $P_1^0 = \{x_i : i = 1,..,n_1\}$, in $P^0$, and where $P_2^0 = P^0 - P_1^0$ denotes the remaining subset of $n_2$ "nonactivated" points in $P^0$. We can then extend the random pattern in Figure S4B to a statistical population of say $N = 999$ random patterns, $P^r$ , $r = 1,..,N$, where each is created by randomly relabeling (permuting) the indices $(1,..,n)$ as $(\pi_{r1},..,\pi_{rn})$, with the initial subset, $P_1^r = \{x_{\pi_{ri}} : i = 1,..,n_1\}$, of $n_1$ points taken to be the permuted version of $P_1^0$ in $P^r$, and with $P_2^r = P^r - P_1^r$ denoting the permuted version of $P_2^0$ . For consistency, all random-permutation tests used here are based on statistical populations of $N = 999$ randomly relabeled patterns. While this sample size may of course be increased, experimentation shows that such increased yield no substantial changes to the results. In these terms, if the observed subpopulation, $P_1^0$, were to exhibit no clustering in $P^0$, the one might expect it to resemble a typical member of to the "null population" of permuted versions, $\{P_1^r : r = 1,..,N\}$.

To test this hypothesis, one must construct an appropriate test statistic for measuring the degree of clustering in each pattern. The most popular measure of clustering is the standard *nearest-neighbor statistic*, which in the present context computes the average distance from each activated cell in a given pattern to its nearest-neighbor activated cell in that pattern. If this average nearest-neighbor distance is smaller for activated cells in the observed pattern, $P^0$, than for say 95% of the random patterns, $P^r$, then this may be said to provide statistically significant evidence of clustering among activated cells in $P^0$.

However, our investigations show that this standard nearest-neighbor procedure exhibits some serious shortcomings in the present application. In essence, this procedure tends to be too "myopic" in the sense that only closeness of nearest neighbors is considered. The present example in Figure S4A provides a good illustration of this shortcoming. For even though there is evident clustering of these activated cells, the standard nearest-neighbor test above shows that there is no significant nearest-neighbor clustering of these cells, and in fact that the average nearest-neighbor distance in pattern, $P^0$, is actually larger than 73% of the $N = 999$ random patterns generated. While the exact reasons for this are not fully known at present, one can speculate that activated cells exhibit local behavior (possibly local movement patterns) that tend to repel nearby cells. In any case, as we shall see below, the clustering behavior of these cells is only evident at slightly larger scales.

What is needed to quantify these scale effects is a test statistic that can actually measure clustering effects at different scales. This is the fundamental property exhibited by *K-function statistics,* which we now develop. In particular, if for any distance value *k*, one counts the number of activated cells within $k\,\pi m$ of any given activated cell, then the average of these counts is designated as the *K-function statistic* at *scale k.* Intuitively, if the average count of activated cells within $k\,\mu m$ of each other is larger than would be expected for typical random permutations of cell locations in pattern, $P^0$, then this suggests some degree of clustering among such cells *at scale k.*

To formalize K-functions themselves, we start with a given pattern, $P^\tau$, and for each target cell, $x_i \in P_1^\tau \subset P^\tau$, let the number of other target cells, $x \in P_1^\tau - \{x_i\}$ within distance *k* of $x_j$ be denoted by,

(1) $$N_k(x_i \mid P_1^\tau) = \#\{x \in P_1^\tau - \{x_i\} : \| x_i - x \| \le k\}$$

In terms of these counts, the *K-function statistic*, $S_\tau(k)$, for pattern, $P^\tau$, at scale *k* is then given by the average value,

(2) $$S_\tau(k) = \tfrac{1}{n_1} \sum_{i=1}^{n_1} N_k(x_i \mid P_1^\tau)$$

The standard *K-function* statistic typically involves a normalization by the spatial density of points in the region of interest, but in the present setting of permutation tests such density considerations play no substantive role.

While this K-function is in principle defined for all distances, $k$, one typically identifies a reasonable range of distance (or scale) values for testing purposes. In the present case, we have chosen the $\mu m$ values, $D = \{1, 2, .., 20\}$. In these terms, the testing procedure for clustering of activated cells at each scale, $k \in D$, can be summarized as follows:

(i) Given observed pattern, $P^0 = \{x_i : i = 1,..,n\}$, with target subset, $P_1^0 = \{x_i : i = 1,..,n_1\}$, generate $N = 999$ random permutations, $(\pi_{ri} : i = 1,..,n)$, $r = 1,..,N$, of $(1,..,n)$ and form the randomly-permuted patterns, $P^r = \{x_{\pi_{ri}} : i = 1,..,n\}$ with corresponding target subsets, $P_1^r = \{x_{\pi_{ri}} : i = 1,..,n_1\}$.

(ii) For each scale, $k \in D$, use expression (2) to compute the observed test statistic, $S_0(k)$, together with corresponding statistics, $S_r(k)$, for each simulated pattern, $P^r, r = 1,..,N$. The appropriate *null hypothesis*, $H_0^S(k)$, to be tested at scale, $k$, is then that *the observed value*, $S_0(k)$, *is drawn from the same statistical population as the set of simulated values*, $\{S_r(k) : r = 1,..,N\}$.

(iii) If we now denote the number of simulated values, $S_r(k)$, at least as large as the observed value, $S_0(k)$, by $m_0^S(k) = \#\{r = 1,..,N : S_r(k) \geq S_0(k)\}$, then the ratio

$$(3) \qquad p_0^S(k) = \frac{m_0^S(k) + 1}{N + 1}$$

denotes the probability of drawing a value as large as $S_0(k)$ from this statistical population of $N+1$ random samples (where "1" corresponds to the observed value treated as an additional random sample). In these terms, $p_0^S(k)$ constitutes the *p-value* for a one-sided test of hypothesis, $H_0^S(k)$. If this p-value is sufficiently small, say $p_0^S(k) \leq 0.05$, then one may infer significant *clustering* of the target population, $P_1^0$, at scale *k*. Conversely if this p-value is sufficiently large, say $p_0^S(k) \geq 0.95$, the one may infer significant *dispersion* of the target population, $P_1^0$, at scale *k*.

The results of these tests for each $k \in D$ are plotted in Figure S5 for the observed pattern, $P^0$, in Figure S4A. Here it can be seen that there is very significant clustering of activated cells beyond 3 $\mu m$. Note also that the number of activated pairs in $P^0$ separated by no more than 1 $\mu m$ is only about the median value for randomly permuted patterns. This reflects the observation above that average nearest-neighbor distances among activated cells in are actually larger than for most randomly permuted patterns. Thus, the significant clustering among such cells at all scales beyond 3 $\mu m$ would be completely missed by tests based on nearest neighbors.

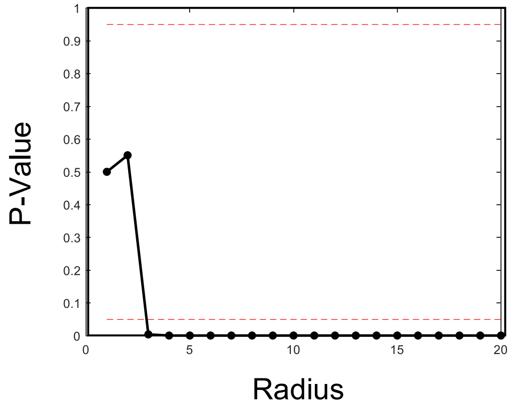


**Figure S5. K-function statistical analysis of clustering.** Clustering of Nur77$^{GFP+}$ OT-I T cells compared to Nur77$^{GFP-}$ OT-I T cells showing significant clustering at a radius (*k*) of 3 μm.

Tests using Cross K-Functions

Our second procedure is again best illustrated by a concrete example. Figure 3K in the text focuses on the spatial relations between uninfected cDC1 cells and activated Nur77$^{GFP+}$cells during early T cell priming events, and displays test results for six milky spots in the omenta of two *snx22*$^{GFP/+}$mice. Figure S6A shows the observed cell pattern for one of these milky spots, containing $n_1 = 261$ Nur77$^{GFP+}$cells (yellow) and $n_2 = 1270$ cDC1 cells (green). Of specific interest is whether the green cDC1 cells tend to cluster around the yellow Nur77$^{GFP+}$ cells more than would be expected in random arrangements of these cells. One such random arrangement is shown in Figure S6B, and suggests that there is indeed some degree of clustering. Notice for example that in the sparser left half of each figure the random arrangement on the right contains many more yellow cells with no green cells close by.

To quantity such associations at different scales, we again adopt a K-function approach. But here we focus on "cross" counts between populations. More specifically, for any given separation distance, *k*, we ask whether the average number of green cells within distance *k* of any yellow cell is larger than would be expected for a random arrangement of these cells. If so, then this will be said to provide evidence for *clustering* of green cells around yellow cells at *scale k*. In the literature, such relations between separate populations are often referred to as "attraction-repulsion" relations, to distinguish them from "cluster-dispersion" relations within a single population. To avoid any causal implications here, we choose to stay with the more descriptive "cluster-dispersion" terminology.

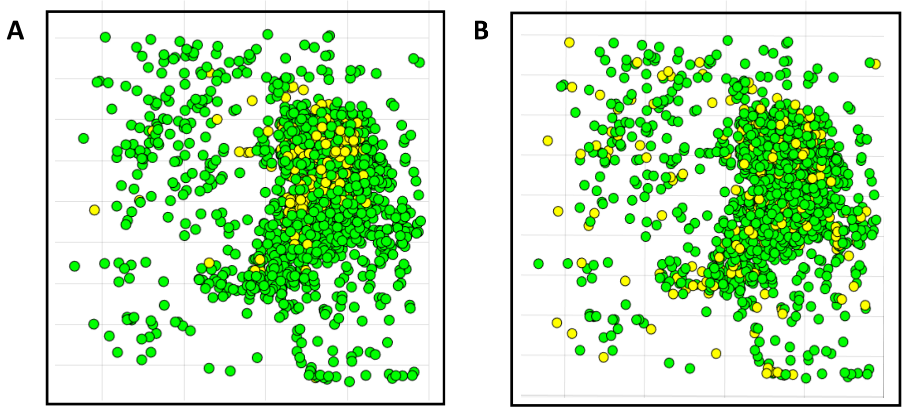


**Figure S6. Cross K-function statistical analysis of T cell clustering.**
(A-B) Intraperitoneal transfer of 5 x 10$^5$ CTV-labeled OT-I/Nur77$^{GFP}$ T cells into *snx22*$^{GFP/+}$ mice immunized with 2 x 10$^5$ CPS-OVA parasites 2 hr later. Immunofluorescent images were analyzed to determine the coordinates of OT-I T cells and cDC1s. (A) 2-dimensional (2D) model representation of coordinates of Nur77$^{GFP+}$ OT-I T cells (yellow) and cDC1s (green) in a MS.
(B) Representative randomized distribution of Nur77$^{GFP+}$ OT-I T cells and cDC1s from (A).

This is easily formalized by an appropriate modification of the K-function definitions above. We again start with a cell pattern, $P^\tau = (x_i : i = 1,..,n)$, consisting of two types of cells: a "target" subpopulation of *type 1* cells, $P_1^\tau = (x_i : i = 1,..,n_1)$, and "reference" subpopulation of *type 2* cells, $P_2^\tau = (x_j : j = n_1 + 1,..,n_1 + n_2 = n)$, where we now let $x_i$ and $x_j$ denote typical elements of $P_1^\tau$ and $P_2^\tau$, respectively. In the present example, types 1 and 2 correspond, respectively, to yellow (Nur77$^{GFP+}$) cells and green (cDC1) cells. More generally, if for each type 1 cell, $x_i \in P_1^\tau \subset P^\tau$, we now let the number of type 2 cells, $x_j \in P_2^\tau$, located within distance *k* of $x_i$ be denoted by,

(4) $$N_k(x_i \mid P_2^\tau) = \#\{x_j \in P_2^\tau : \| x_i - x_j \| \leq k\} \ , \ x_i \in P_1^\tau$$

then the *cross K-function statistic*, $C_\tau(k)$, for pattern, $P^\tau$, at *scale k* is given by the average of these count values,

(5) $$C_\tau(k) = \tfrac{1}{n_1} \sum_{i=1}^{n_1} N_k(x_i \mid P_2^\tau)$$

We again drop any reference to point densities in defining these *cross K-function* statistics for purposes of random permutation tests.

With these definitions, the testing procedure for identifying clustering of type 2 cells around type 1 cells at each scale, $k \in D = \{1,2,..,20\}$, amounts essentially to replacing K-function statistics, $S(k)$, with cross K-function statistics, $C(k)$. For completeness, this test is summarized as follows:

(i) Given observed pattern, $P^0 = \{x_i : i = 1,..,n\}$, with target subset, $P_1^0 = \{x_i : i = 1,..,n_1\}$ and reference subset, $P_2^0 = \{x_j : j = n_1+1,..,n_1+n_2 = n\}$, generate $N$ = 999 random permutations, $(\pi_{ri} : i = 1,..,n)$, $r = 1,..,N$, of $(1,..,n)$ and form the randomly-permuted patterns, $P^r = \{x_{\pi_{ri}} : i = 1,..,n\}$ with corresponding subsets of type 1 cells, $P_1^r = \{x_{\pi_{ri}} : i = 1,..,n_1\}$, and type 2 cells, $P_2^r = \{x_{\pi_{rj}} : j = n_1+1,..,n\}$.
(ii) For each scale, $k \in D$, use expression (5) to compute the observed test statistic, $C_0(k)$, together with corresponding statistics, $C_r(k)$, for each simulated pattern, $P^r, r = 1,..,N$. The appropriate *null hypothesis*, $H_0^C(k)$, to be tested at scale, $k$, is then that *the observed value*, $C_0(k)$, *is drawn from the same statistical population as the set of simulated values*, $\{C_r(k) : r = 1,..,N\}$.
(iii) If we now denote the number of simulated values, $C_r(k)$, at least as large as the observed value, $C_0(k)$, by $m_0^C(k) = \#\{r = 1,..,N : C_r(k) \geq C_0(k)\}$, then [as in (3) above] the ratio,

(6) $$p_0^C(k) = \frac{m_0^C(k)+1}{N+1}$$

now constitutes the *p-value* for a one-sided test of hypothesis, $H_0^C(k)$. If this p-value is sufficiently small, say $p_0^C(k) \leq 0.05$, then one may infer significant *clustering* of type 2 cells around type 1 cells in pattern $P^0$ at scale *k*. Conversely if $p_0^C(k)$ is sufficiently large, say $p_0^C(k) \geq 0.95$, then one may infer significant *dispersion* of type 2 cells around type 1 cells in pattern $P^0$ at scale *k*.

The results of these tests for each $k \in D$ are plotted in Figure S7 below for the observed pattern in Figure S6A above. In Figure 3K of the text, this plot corresponds to the unique plot starting at p-value = 0.65. Here it can be seen that there is very significant clustering of cDC1 cells around Nur77$^{GFP+}$cells starting at 4 $\mu m$. Note finally that all test results summarized in Figure 3K indicate similar significant clustering starting at around 5 $\mu m$. However, this degree of significance is seen to be more variable at scales beyond 8 $\mu m$, suggesting that clustering behavior between these cell populations may be limited in scale.

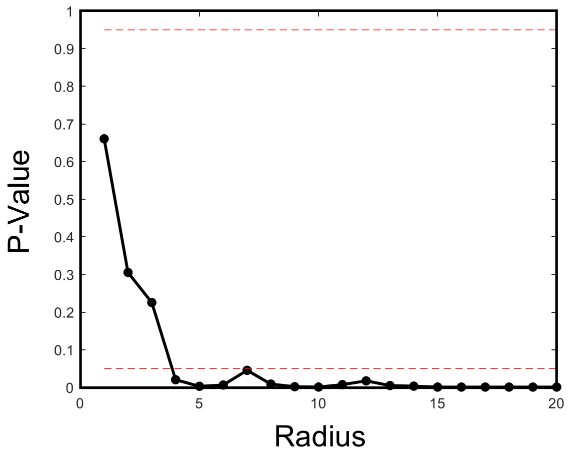


**Figure S7. Cross K-function statistical analysis of clustering.** Analysis of cDC1 clustering around Nur77$^{GFP+}$ OT-I T cells showing significant clustering at a radius (*k*) of 4 μm.
